# A cleaved cytosolic FOXG1 promotes excitatory neurogenesis by enhancing mitochondrial translation

**DOI:** 10.1101/2024.08.13.607559

**Authors:** Hannah Bruce, Kathryn Adamson, Gessica Goncalves, Marnie E. Halpern, Clinton Monfries, Oniz Suleyman, Lourdes Osuna-Almagro, Fursham Hamid, Katherine R. Long, Corinne Houart

**Affiliations:** MRC Centre for Neurodevelopmental Disorders, Institute of Psychiatry, Psychology & Neuroscience, King’s College London, London, SE1 1UL; Centre for Developmental Neurobiology, Institute of Psychiatry, Psychology & Neuroscience, King’s College London, London, SE1 1UL; Francis Crick Institute, Midland Road, London, NW1 1AT; Geisel School of Medicine, Molecular and Systems Biology, Dartmouth, Hanover, NH03755

## Abstract

The modulation of mitochondrial function is core to cell fate decisions and tissue homeostasis, yet the mechanisms regulating these processes remain poorly understood. Here, we demonstrate that mitochondrial OXPHOS activity is controlled in a tissue-specific manner through a non-canonical cytoplasmic function of the transcription factor FOXG1. A temporally regulated short cytoplasmic FOXG1 interacts with mitochondrial ribosomal proteins and enhances translation of mt-DNA-encoded OXPHOS proteins. Zebrafish and human models of early nonsense FOXG1 syndrome mutations unexpectedly produce a short C-terminal peptide. Expression of this truncated protein drives an overproduction of excitatory neurons and induces a structural, functional, and translational phenotype in mutant mitochondria. We demonstrate that this activity is a gain of function, normally carried out by a cleaved FOXG1 in wildtype. Both normal and mutant peptides are transported to the mitochondria, interact with mito-ribosomal proteins to enhance translation, thereby stimulating neurogenesis. Adjusting the dosage of the mutant peptide rescues the excitatory aspect of the FOXG1 syndrome. Our study demonstrates a novel role for cytoplasmic Foxg1 in promoting neurogenesis via the tuning of mitochondrial translation and provides the first evidence for direct tissue-specific control of OXPHOS activity. This study opens a novel therapeutic avenue for the treatment of disorders associated with mitochondrial hypoactivity.

**Graphic Abstract:** 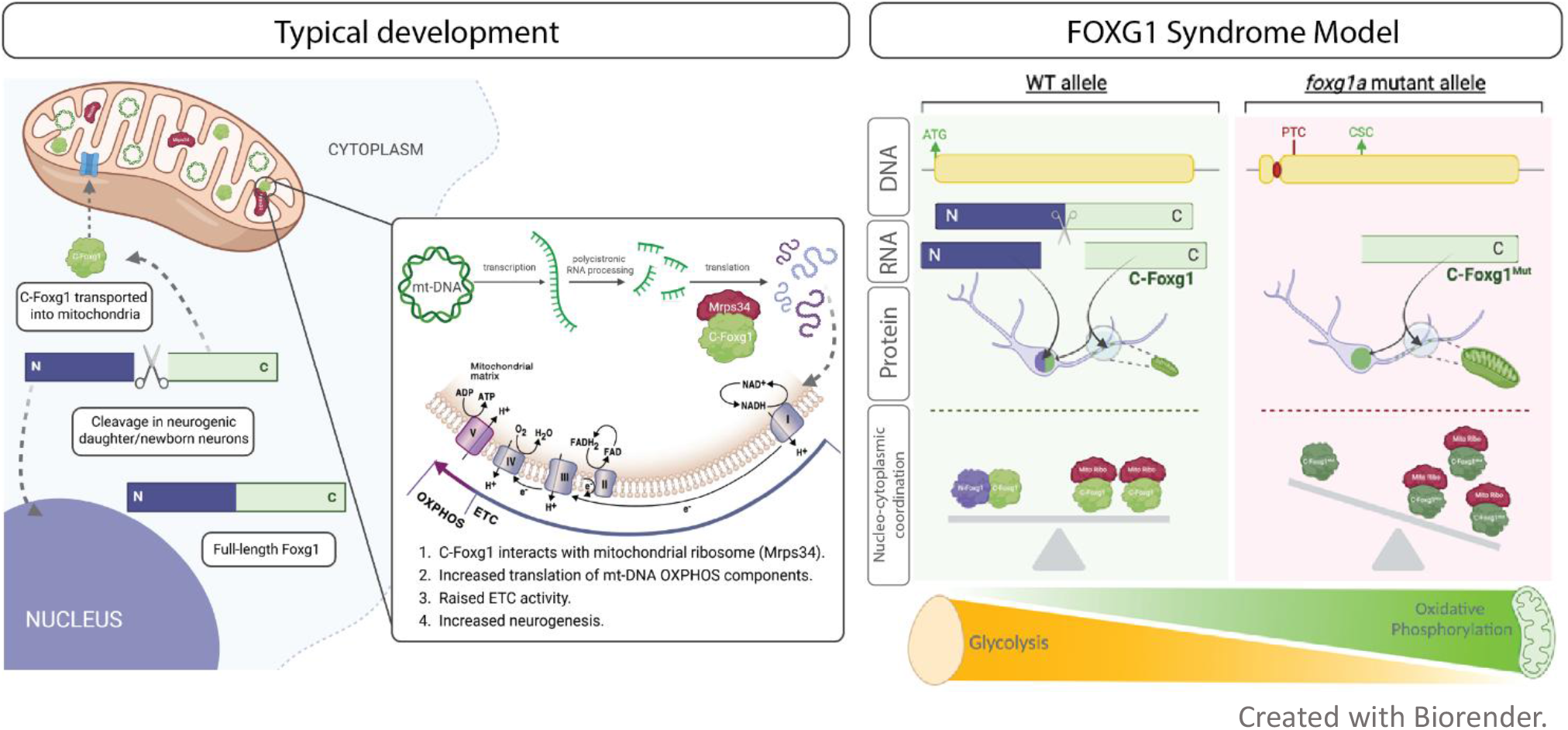

## Introduction

Mitochondria are pivotal regulators of cell fate decisions and have recently been identified as pacemakers in relation to species-specific development ^1^. In the brain, multiple studies have demonstrated that energy production shifts during differentiation: stem cells depend primarily on glycolysis to produce ATP, whereas cells transitioning to neurons exhibit higher energy demand and increasingly rely on oxidative phosphorylation (OXPHOS)^2,3^. Stimulating OXPHOS accelerates neuronal differentiation, underscoring the organelle’s role in pacing cell fate decisions ^1,4^. Yet, the mechanisms that regulate mitochondrial activity during these shifts remains poorly understood.

Models of neurodevelopmental disease offer a valuable framework for the study of human brain development. FOXG1 Syndrome (*OMIM: 613454*) is one such disorder, caused by heterozygous disruptions in the Forkhead Box G1 (*FOXG1*) gene, which encodes a transcription factor essential for development of the telencephalon^5^. It is predominantly characterised by microcephaly, corpus callosum (CC) agenesis, and seizures^6^. FOXG1 dysregulation is also implicated in autism spectrum disorder (ASD) and schizophrenia ^7,8^. FOXG1 is amongst the earliest transcription factors to be induced in the prospective forebrain and is highly expressed in telencephalic progenitors. Homozygous deletion of *Foxg1* in mice^5^, or *foxg1a* KD in zebrafish^9^, leads to a drastic reduction in the size of the telencephalon as a result of premature neuronal differentiation, and depletion of the progenitor pool. Complete loss of Foxg1 results in a failure to specify the ventral telencephalon, subsequent dorsalization of the tissue^10^, and a nearly complete loss of inhibitory interneurons (IN). Reduction of IN numbers is also observed in human FOXG1 Syndrome post-mortem cortical samples^10–12^. Most studies investigating FOXG1 Syndrome mechanisms have focused on *null* mutations, which do not accurately model the human heterozygous condition. However more recently heterozygous murine models have been revisited, and advances in technology have revealed subtle but important phenotypes, including proliferation deficits, axonal guidance impairment, and disruption of cellular excitatory/inhibitory (E/I) balance^15–17^.

FOXG1’s roles in proliferation and neuronal differentiation may be segregated across subcellular compartments. Progenitors display a predominantly nuclear localisation of FOXG1, while differentiating cells exhibit some additional cytoplasmic localisation^13^. In *Xenopus*, post-translational modifications of FOXG1 influence the localisation of the protein: phosphorylation of a serine residue (Ser 19) promotes nuclear import, while phosphorylation of a threonine (Thr 226) in the C-terminus enhances nuclear export ^13^. In mouse neuronal culture, the protein can undergo post-translational cleavage to produce a C-terminal fragment (C-FOXG1) that co-localizes with mitochondria and an N-terminal product that displays an exclusively nuclear localisation ^14^. Our study provides answers to the unsolved physiological functions of FOXG1 cytoplasmic localisation and cleavage products.

To investigate the role of non-nuclear Foxg1 and the temporal pattern of its expression in a disease context, we generated a heterozygous *foxg1a* early frameshift zebrafish mutant. The line harbours a 5-basepair (bp) deletion close to the canonical start codon, leading to the induction of a premature termination codon (PTC, *foxg1a^+/5bp^*). Importantly, PTCs represent a third of all FOXG1 Syndrome patient genetic lesions and are associated with the most severe clinical presentations, predicted to result in a *null* allele phenotype^6^. We demonstrate that *foxg1a^+/5bp^* mutants recapitulate key features of the human condition and challenge the prevailing view that early PTC-generating mutations abolish protein expression. Instead, we show that the zebrafish mutant mRNA adopts a downstream in-frame ATG as a cryptic start codon (CSC), translation from which produces a C-terminal mutant Foxg1 peptide (C-Foxg1^Mut^). We found that this mechanism is conserved in human iPSC-derived neuronal cells harbouring a *FOXG1* patient mutation. Functional analyses in zebrafish revealed that C-Foxg1^Mut^ function significantly contributes to the brain pathology in this model. Our study reveals that C-Foxg1^Mut^ is a gain-of-function of the normal cleavage product, C-Foxg1. Both peptides are found in the cytoplasm of human, mouse, and zebrafish newborn neurons, more abundantly in excitatory cells, and often in mitochondria. Zebrafish *foxg1a^+/5bp^* mutants display a pronounced increase in the expression of mitochondrial ribosomal and mt-DNA encoded OXPHOS proteins, a robust C-Foxg1^Mut^-dependent hyperactive mitochondrial phenotype, which leads to an excess in excitatory neuronal numbers. Co-immunoprecipitation experiments confirm a direct interaction of C-Foxg1 with mito-ribosomal proteins, further supporting a role for C-Foxg1 in enhancing mitochondrial translation.

Overall, our study demonstrates a novel role for cytoplasmic Foxg1 in promoting neurogenesis via the tuning of mitochondrial translation, providing the first evidence for direct tissue-specific instructive control of OXPHOS activity. We uncover a pathogenic gain of this mitochondrial function in fish and human disease models, unveiling a previously unknown therapeutic target and opening a potential therapeutic avenue for the treatment of many disorders involving mitochondrial tuning dysfunctions.

## Results

Antibody staining in zebrafish (Figure 1A) and human foetal telencephalon (Figure 1B) revealed a pool of non-nuclear Foxg1, excluded from neuroepithelial progenitors. Cytoplasmic Foxg1 is found in neuronal cells, partly colocalizing with mitochondrial markers (COX-IV; fish, TOMM20; human). To determine if non-nuclear Foxg1 was inside the mitochondria, we performed expansion microscopy on 48hpf zebrafish embryos and focused our image acquisition on the dorsolateral telencephalon to include predominantly neurons. A representative 3D projection can be seen in Figure 1C, demonstrating that Foxg1 does indeed exhibit mitochondrial localization. These observations led us to further investigate the role of Foxg1 in the regulation of mitochondrial function.

**Figure 1:**
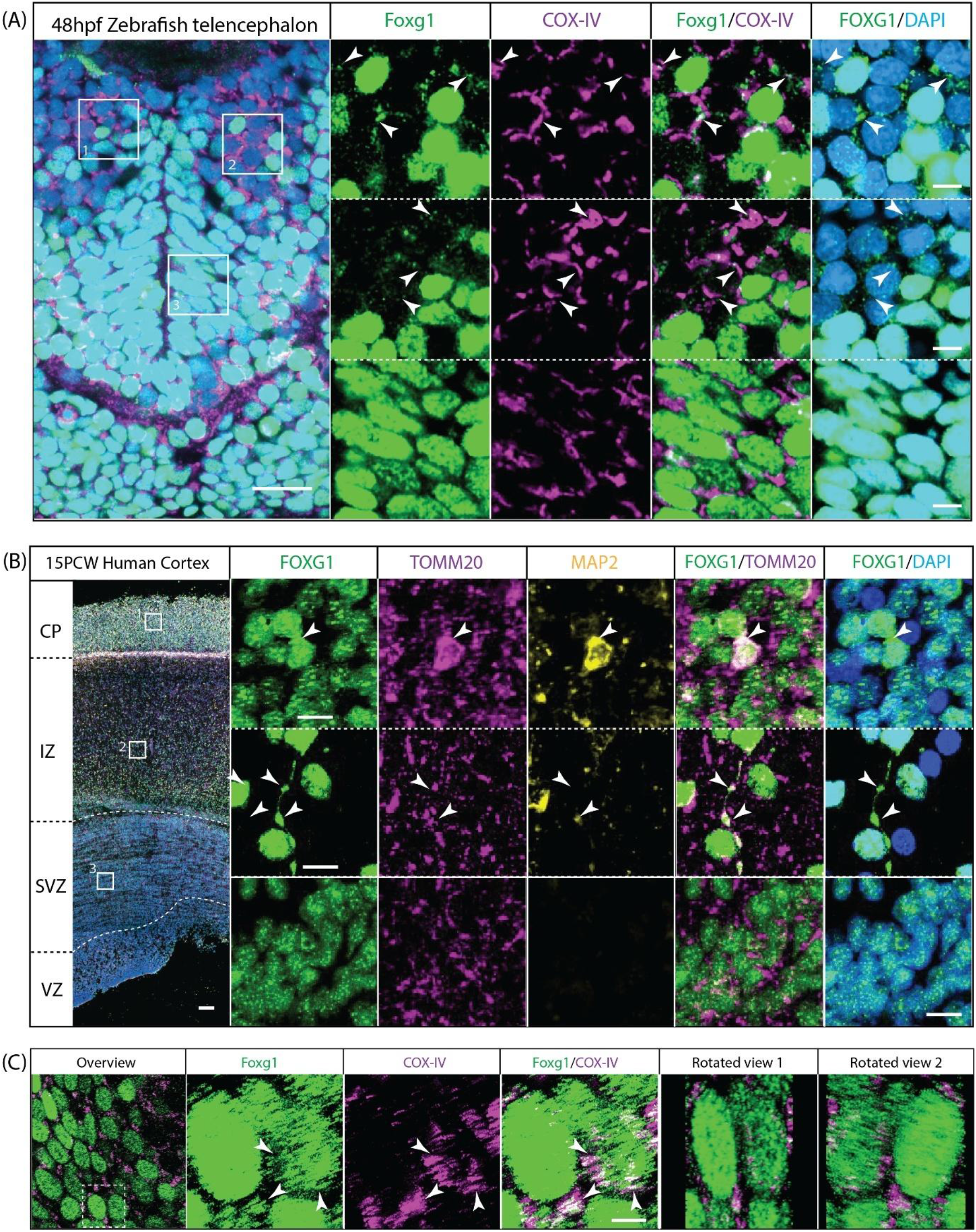
Non-nuclear Foxg1 is abundant in neurons but not progenitors. **A.** 48hpf wild type zebrafish telencephalon stained for Foxg1 (green), COX-IV (magenta), and DAPI (blue); images are single planes. Scale bars = 20µm (overview) and 5 µm (insets). **B.** 15 post conception weeks (PCW) human foetal telencephalon stained for FOXG1 (green), TOMM20 (magenta), MAP2 (yellow), and DAPI (blue); image shown are projections of small z-stacks. Scale bars = 100µm (overview) and 5 µm (insets). **C.** Expansion microscopy of Foxg1 (green) and COX-IV (magenta) stained 48hpf wild type zebrafish telencephalon. Overview image is a single slice of a z-stack, while inset images are 3D projections. Two 3D-rotated views of the projection can be seen in the final two panels. Scale bar = 5µm.

### Generating a zebrafish model of FOXG1 Syndrome to investigate the function of non-nuclear Foxg1

We generated a zebrafish model of the most common mutations reported in FOXG1 Syndrome: early frameshift mutations. Deletion of 5bp close to the canonical start codon of *foxg1a* leads to the induction of a premature termination codon (PTC), with no protein predicted to be produced. To validate the *foxg1a^+/5bp^* zebrafish model we used a complex transgenic line: (*Tg(eomesa:QF2)^c7^*^14^ *;Tg(QUAS:mApple-CAAX; he1.1:mCherry)^c6^*^36^ and *Tg(Dlx5a-6a;GFP*)), hereafter referred to as ‘double transgenic’. This line allows endogenous mCherry membrane labelling of excitatory neurons (Eomesa^+^, aka Tbr2^+^), while inhibitory interneurons (Dlx^+^) are identified by GFP whole cell labelling. At 28hpf and 48hpf the *foxg1a^+/5bp^* mutant exhibits a significant decrease in the number of inhibitory neurons and increase in the number of excitatory neurons, leading to a shift in cellular E/I balance toward excitation (Figure 2A). The possibility of compensation for this imbalance at the synapse level was ruled out by mosaic labelling of excitatory synapses via QUAS-dependent expression of synaptophysin (Supplementary Fig. 1A). Unlike mammals, the teleost telencephalon lacks a CC, with the anterior commissure (AC) being the prominent cross-hemisphere connection. At 28hpf there was no significant difference in the thickness, width, or volume of the AC in *foxg1a^+/5bp^* compared to *foxg1a^+/+^*, indicating no developmental delay in axonal outgrowth (Figure 2B). However, by 48hpf the thickness and volume of the AC was significantly decreased in heterozygous mutants (p = 0.018, unpaired t test; p = 0.0078, unpaired t test). To assess the progenitor/neuron balance in *foxg1a^+/5bp^* mutants we used a transgenic line, *Tg(HuC:GFP)*, that expresses GFP under control of the early neuronal marker ELAV3/HuC promoter and examined HuC^+^ versus HuC^-^ cells in the telencephalon^18^. Compared to wildtype, we found an early increase in numbers of post-mitotic neurons (HuC^+^), and a decreased number of progenitors (HuC^-^) in heterozygous mutants, while at later stages both cell types were significantly decreased in *foxg1a^+/5bp^* (Supplementary Fig. 1E). We performed bulk RNA-sequencing at 24hpf to characterise the mutant transcriptome and found significant changes in the expression of genes with annotated functions in neuronal development, cell adhesion, and ATP generation (Supplementary Fig. 2). Collectively, these experiments establish the validity of our heterozygous zebrafish model, demonstrating a cellular E/I imbalance reminiscent of patient seizures, an AC phenotype that recapitulates CC agenesis, and progenitor pool depletion that underlies microcephaly.

**Figure 2:**
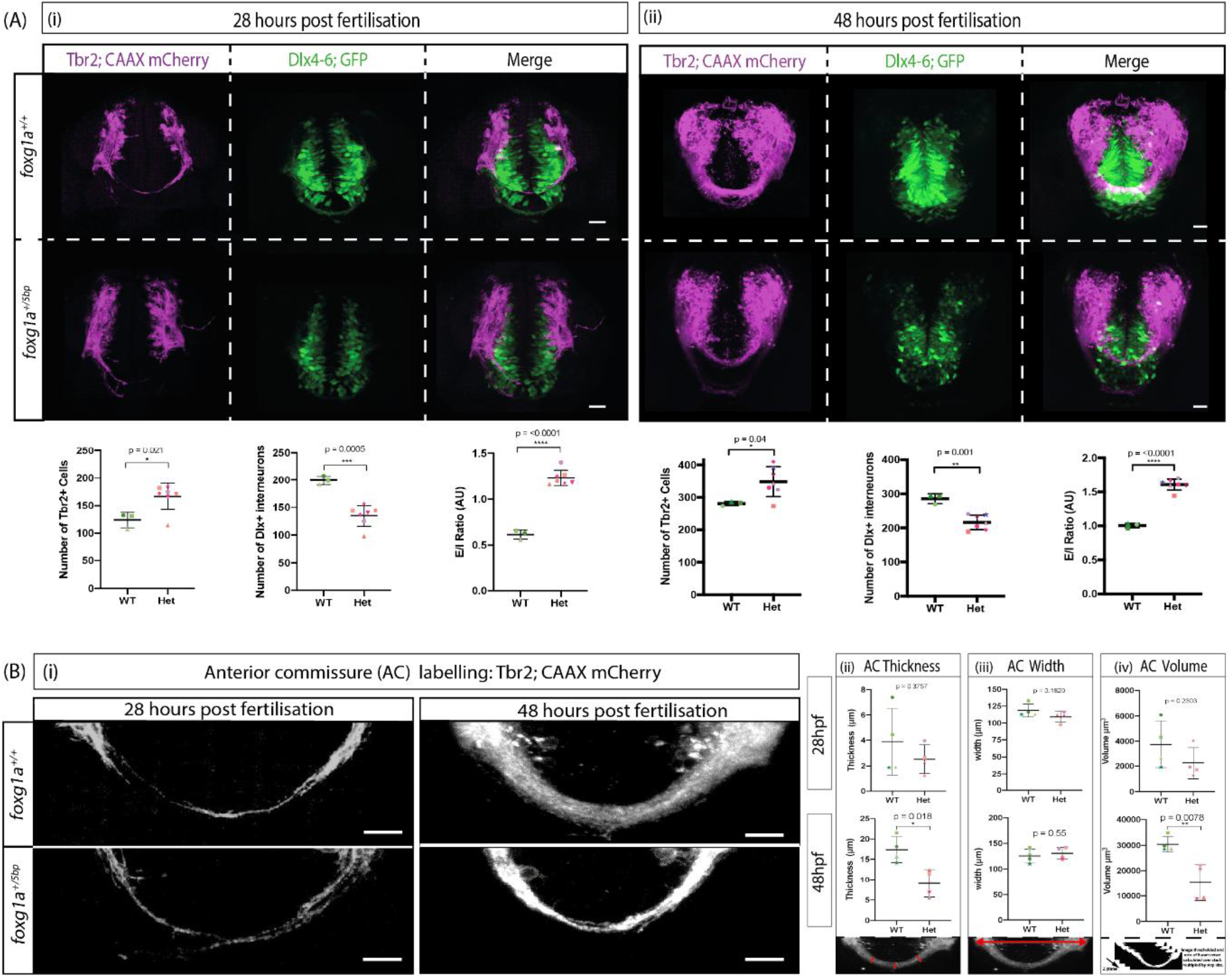
Phenotypic characterization of heterozygous foxg1a+/5bp mutants. A. Representative images are projections of confocal stacks taken in a frontal view; dorsal to the top, at 28hpf (i) and 48hpf (ii) p values calculated using unpaired t tests; for both stages n = 3 *foxg1a^+/+^, 7 foxg1a^+/5pb^*. Scale bars = 20µm; error bars indicate Standard Deviation from the Mean (SDM); statistics from unpaired t tests. B. Representative images are projections of confocal stacks taken in a frontal view; dorsal to the top, at 28hpf and 48hpf. For both stages and genotype n = 4, p value calculated from unpaired t-test. Schematics under charts indicated how the measurement was performed. Scale bars = 20µm; error bars indicate SDM; statistics from unpaired t tests.

### Early frameshift FOXG1 Syndrome mutations produce a cytoplasm-enriched Foxg1 C-terminal peptide that promotes excitatory neurogenesis

Immunostaining with a C-terminal Foxg1 antibody in the *foxg1a^5bp/5bp^* mutant unexpectedly revealed detectable protein expression in the telencephalon and nasal retina. The signal was weaker than in siblings and exhibited an altered subcellular localisation (weak nuclear and abundant cytoplasmic) compared to *foxg1a^+/+^* (Supplementary Fig. 3A). Deep sequencing of *foxg1a^5bp/5bp^* whole embryo transcriptomes (Supplementary Fig. 2) did not support genetic compensation as an explanation for these observations. Instead, Western Blots using antibodies raised against the N or C terminus of the Foxg1 protein showed that *foxg1a* mutants exhibited a smaller band of 35kDa, representing a truncated C-terminal Foxg1 peptide (C-Foxg1^Mut^) (Figure 3A). With the N-terminal antibody (Figure 3A ii) the *foxg1a^+/+^* and *foxg1a^+/5bp^* fish displayed bands at 50kDa (full-length Foxg1) and 30kDa (N-terminal post-translational cleavage product, confirming findings in mice^14^), while the homozygous mutant showed no band. With the C-terminal antibody, *foxg1a^+/+^* and *foxg1a^+/5bp^* fish displayed bands corresponding to full-length Foxg1 alongside a smaller 25kDa cleavage band. This fragment has previously been shown to localise to the mitochondria in mouse cell culture misexpression experiments^19^. The 35kDa C-Foxg1^Mut^ band is consistent with re-initiation of translation from the first ATG downstream of the PTC (herein termed ATG4&5). Antibody staining on homozygous embryos carrying a C-terminal deletion (*foxg1a^ΔC-term/ΔC-term^*, Methods) produced no immunoreactivity, validating the band as a product of translation from CSC (Supplementary Fig. 4C).

**Figure 3:**
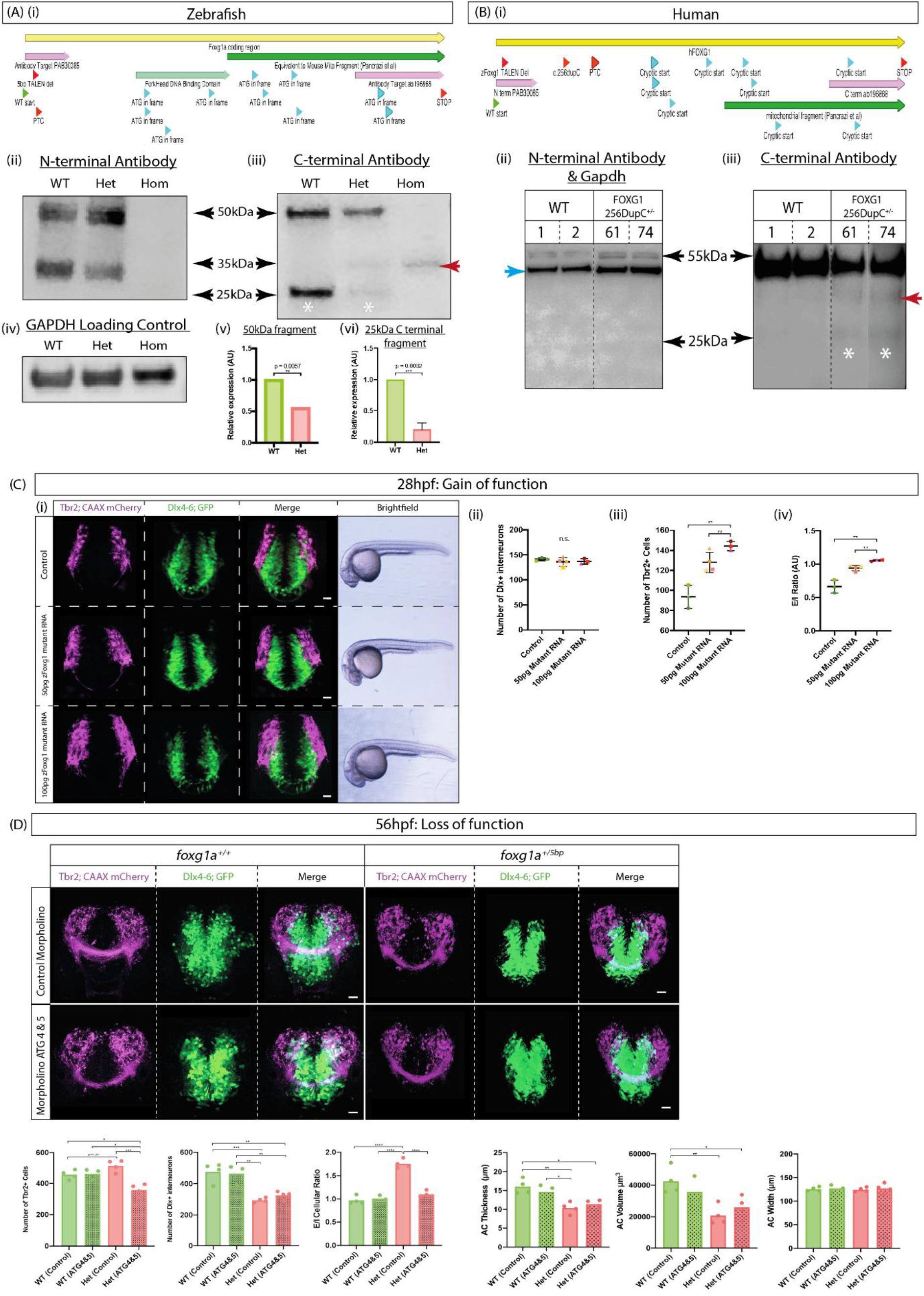
Re-initiation of translation from a cryptic start codon produces a variant C-terminal Foxg1. (A) Zebrafish mutation and protein expression. (i) Schematic of the zebrafish *foxg1a* gene showing 5bp TALEN deletion which causes a frameshift and the induction of a premature termination codon (PTC); further downstream are 10 ATGs in-frame with the canonical stop codon. (ii) Western Blot on 24hpf total lysate with antibody raised against the N-terminal of Foxg1. *foxg1a^+/+^* and *foxg1a^+/5bp^* show a full-length band of ∼50kDa and a smaller ∼30kDa; the *foxg1a^5bp/5bp^* mutant lacks any bands. (iii) Western Blot on 24hpf total lysate with antibody raised against the C-terminal of Foxg1. *foxg1a^+/+^* and *foxg1a^+/5bp^* show a full-length band of ∼50kDa and a smaller ∼25kDa; the *foxg1a^+/5bp^* and *foxg1a^5bp/5bp^* lysate has an additional band at ∼35kDa. (iv) Western Blot on 24hpf total lysate with antibody raised against Gapdh as a loading control; 50ug protein loaded per well. (v) Quantification of the relative expression of full-length (∼50kDa) Foxg1, normalized to Gapdh expression, error bars indicate SDM (n =3). (vi) Quantification of the relative expression of wildtype cleaved C-Foxg1, normalized to Gapdh expression, error bars indicate SDM (n =3). (B) Human mutation and protein expression. (i) Schematic of the human *FOXG1* gene showing the patient frameshift mutation *256 duplication C*, predicted to lead to PTC; further downstream are 9 ATGs in-frame with the canonical stop codon. (ii) Western Blot on DIV 20 neuronal progenitor cells derived from two isogenic iPSC line clones (controls and *FOXG1^256DupC+/-^* mutants) with N-terminal FOXG1 antibody and GAPDH (blue arrow) as a loading control. Both control and *FOXG1 ^+/256DupC^* exhibit a full-length FOXG1 band at ∼55kDa and GAPDH at ∼37kDa. Please note an irrelevant lane was removed from between *FOXG1^+/256DupC^* Clones 61 & 74. (iii) Western Blot on DIV 20 neuronal progenitor cells derived from iPSCs, with the C-terminal antibody. WT and *FOXG1^+/256DupC^* exhibit a full-length FOXG1 band at ∼55kDa and a faint smaller band at ∼25kDa. *FOXG1^+/256DupC^* and a band at ∼35kDa; 50ug of protein per well. Please note an irrelevant lane was removed from between *FOXG1 ^+/256DupC^* Clones 61 & 74. (C) Gain of function - Overexpression of mutant mRNA transcript in WT fish. (i) Representative images of 28hpf controls, 50pg mutant mRNA, and 100pg mutant mRNA shown in a frontal view; scale bars= 20µm. n = 3; uninjected, 4; 50pg, 3; 100pg. (ii) Number of Dlx^+^ interneurons; no significant difference (One-way ANOVA). (iii) Number of Tbr2^+^ excitatory neurons; significantly increased in injected embryos *p = 0.0049* (50pg) and *p = 0.0008* (100pg), One-way ANOVA. (iv) Excitatory/inhibitory (E/I) cellular ratio; *p= 0.001* (50pg) and *p = 0.0002* (100pg) One-way ANOVA. (D) Loss of function - Inhibition of cryptic start codon by MO injection. (i) Representative images of *foxg1a^+/+^* and *foxg1a^+/5bp^* 56hpf embryos injected with either a control MO or a MO against ATG4&5 (CSCs). Images are shown in a frontal view, scale bars= 20µm. (ii) Quantification of number of Tbr2^+^ cells; Two-way ANOVA. (iii) Quantification of number of Dlx^+^ cells; Two-way ANOVA. (iv) Quantification of E/I cellular ratio; Two-way ANOVA. (v) Quantification of anterior commissure (AC) thickness; Two-way ANOVA. (vi) Quantification of AC volume; Two-way ANOVA. (vii) Quantification of AC width; Two-way ANOVA.

Nonsense and early frameshift mutations in the N-terminus of the *FOXG1* gene are responsible for a third of all FOXG1 Syndrome mutations, giving our findings a strong translational relevance. To ask whether patient-derived telencephalic cells also produce a C-FOXG1^Mut^ peptide, we first obtained iPSCs from a patient harbouring a duplication of cysteine at position 256 (*FOXG1^256DupC+/-^*), causing a frameshift and induction of a PTC (Supplementary Fig. 3B). However, to account for differences in genetic background, we also produced isogenic lines carrying the same mutation in a well-characterised and previously established control iPSC line (036S_CTM)^20^.This specific patient mutation occurs upstream of the candidate CSCs in the zebrafish gene, which are themselves conserved from fish to human (Figure 3Bi). For neuronal patterning of iPSCs, we performed a dual SMAD inhibition^21^ and repeated N & C terminal antibody Western Blots on day *in vitro* 20 (DIV 20) human neuronal progenitor cells (NPCs). Control and mutant NPCs displayed bands corresponding to full-length FOXG1 (∼55kDa; human protein is larger than fish) in both Western conditions, but mutant NPCs also produced a smaller peptide, again corresponding to re-initiation of translation from the first downstream CSC.

Given the conservation of the cryptic ATG, we next assessed whether expression of C-Foxg1^Mut^ has functional consequences. We performed a gain of function (GOF) experiment, injecting varying doses of the mutant *foxg1a* mRNA into wildtype *foxg1a^+/+^* double transgenics (excitatory and inhibitory cell labelling) and quantified numbers of neurons at 28hpf and 56hpf. We observed a robust dose-response increase in the number of Tbr2^+^ excitatory neurons (Figure 3C) and no difference in the number of Dlx^+^ inhibitory cells, indicating that the peptide is preferentially active and/or translated in the pallium. This effect was maintained at later developmental stages (Supplementary Fig. 3C), showing that C-Foxg1^Mut^ is sufficient to increase excitatory neurogenesis.

We then asked whether C-Foxg1^Mut^ was also necessary for this increase, assessing if repression of C-Foxg1^Mut^ activity may improve the mutant phenotype. For this, we injected an antisense morpholino (MO) targeting ATG4&5 which works by preventing translation at the CSC site on the mRNA. Foxg1 antibody staining and Western Blot of 24hpf homozygous *foxg1a^5bp/5bp^* mutants confirmed that levels of protein were substantially decreased upon MO treatment (Supplementary Fig. 3E&F). When C-Foxg1^Mut^ was inhibited in *foxg1a^+/5bp^* mutants at 28hpf (Supplementary Fig. 3D) and 56hpf (Figure 3D), Tbr2^+^ cell numbers were significantly decreased compared to *foxg1a^+/5bp^* fish injected with a control MO (p = 0.0003, Two-way ANOVA). No effect was observed on Dlx*+* inhibitory neurons. The decrease in excitatory cell numbers significantly improved the neuronal E/I ratio in *foxg1a^+/5bp^* embryos. We also assessed the impact of MO treatment on the commissural phenotype at 56hpf and did not observe any improvement. The ATG4&5 MO treatment had no effect on any measurement in wildtype siblings, indicating there was no off-target effect on translation from the canonical ATG of the wildtype allele. All together, these experiments show that in early frameshift heterozygotes, the reduction of inhibitory neurons is due to loss of full-length Foxg1 protein function, while the excess of excitatory neurons is a consequence of gain of function of C-Foxg1^Mut^ expression.

### C-Foxg1^Mut^ localises to the mitochondria and causes a bioenergetic phenotype

Given that the C-Foxg1^Mut^ peptide contains a mitochondrial localizing domain, we postulated that the amount of Foxg1 localising to the mitochondria may be altered in the early frameshift mutant. Using double staining for Foxg1 and a mitochondrial marker (COX-IV), we quantified the subcellular localization of Foxg1 and found that mutant fish exhibited an increase in the percentage of non-nuclear Foxg1, ATG4&5 MO treatment restoring this proportion back to WT levels (Figure 4A v). To quantify how much Foxg1 was localised to the mitochondria, we calculated the percentage of total and non-nuclear Foxg1 protein colocalizing with COX-IV. We found that *foxg1a^+/5bp^* and *foxg1a^5bp/5bp^* mutants show an increase in the percentage of Foxg1 overlapping with COX-IV and that inhibition of CSC function rescued this increase. Using the mitochondria network analysis tool, MiNA^22^, we found that *foxg1a^+/5bp^* and *foxg1^5bp/5bp^* mutants exhibit an increased mitochondrial footprint (number of mitochondria in field of view), and their mitochondria had a significantly increased mean branch length (proxy for mitochondria size, Figure 4A ii). Of note, the difference between wildtype and heterozygous siblings is more pronounced in the pallium, where excitatory neurons are generated. Treatment with the ATG4&5 MO rescues the mitochondrial phenotypes, with no more difference in the mitochondrial footprint or mean branch length observed between wildtype siblings and heterozygous (Figure 4A iv; an example of MiNA segmentation can be found in Supplementary Fig. 4A).

**Figure 4:**
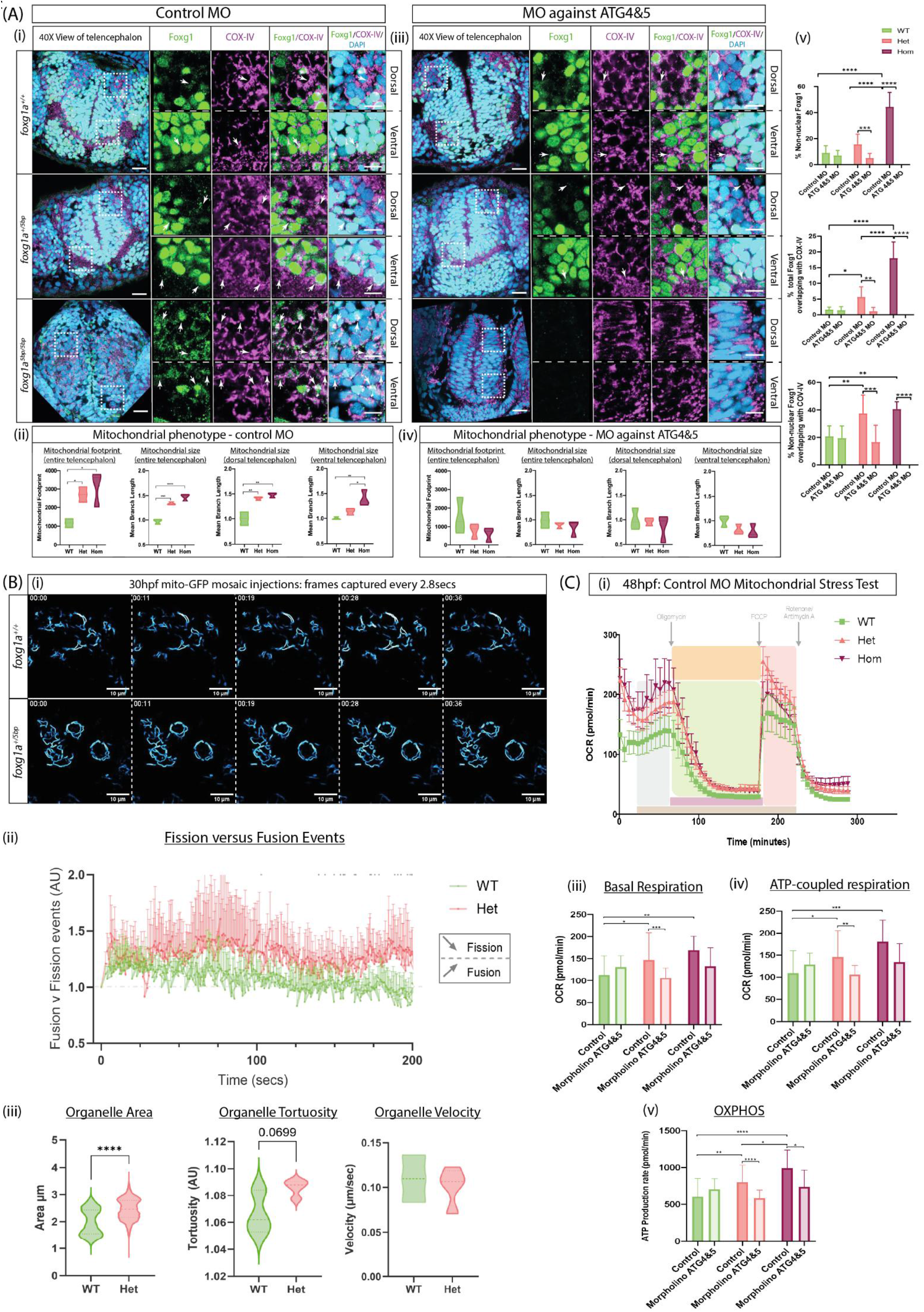
C-Foxg1Mut modifies mitochondrial activity. (A) Mitochondrial phenotype of *foxg1a* mutants (i) Foxg1 and COX-IV antibody staining of 48hpf telencephalon in *foxg1a^+/+^*, *foxg1a^+/5bp^* and *foxg1a^5bp/5bp^* embryos injected with a control MO. Representative images are single slices from a confocal stack taken in a frontal view (insets from the dorsal and ventral telencephalon). Scale bars = 20µm (ii) Mitochondrial footprint (number of mitochondria in field of view) and branch length (size) of *foxg1a^+/5bp^* and *foxg1a^5bp/5bp^* mutants injected with control MO. Mitochondrial footprint was increased in control morpholino treated mutants (*p = 0.005* for *foxg1a^+/5bp^* and *p = 0.026* for *foxg1a^5bp/5bp^*, One way ANOVA). Branch length (mitochondrial size) was also significantly increased across the entire telencephalon in both mutants (*p = 0.0005* for *foxg1a^+/5bp^* and *p = <0.0001* for *foxg1a^5bp/5bp^*, One way ANOVA). MO ATG4&5 rescued mitochondrial phenotypes in mutants. n = 3 *foxg1a^+/+^,* 3 *foxg1a^+/5bp^*, 3 *foxg1a^5bp/5bp^*; ATG4&5 MO n = 3 *foxg1a^+/+^,* 3 *foxg1a^+/5bp^*, 3 *foxg1a^5bp/5bp^*. (iii) Foxg1 and COX-IV antibody staining of 48hpf telencephalon in *foxg1a^+/+^*, *foxg1a^+/5bp^* and *foxg1a^5bp/5bp^* embryos injected with MO against ATG4&5. Representative images are single slices from a confocal stack taken in a frontal view (insets from the dorsal and ventral telencephalon). Scale bars = 20µm (iv) Mitochondrial footprint and branch length of *foxg1a^+/5bp^* and *foxg1a^5bp/5bp^* mutants injected with MO against ATG4&5. n = 3 for each genotype. (v) Mutant zebrafish exhibit an increase in the non-nuclear percentage of Foxg1 and an increase in the percentage of Foxg1 overlapping with the mitochondria (COX-IV). MO treatment had no effect on any measurement in WT fish but rescued the increased percentage of Foxg1 overlapping with the mitochondria in mutants. Statistics calculated with Two-way ANOVAs. Note that ATG4&5 MO effectively reduces protein signal to zero in homozygous mutants. (B) Timelapse imaging of mitochondrial dynamics (i) Representative frames from *foxg1a^+/+^* and *foxg1a^+/5bp^* 30hpf dorsal telencephalon showing mosaic mito-GFP labelling in Tbr2+ neurons. Frames were captured every 2.8 seconds for 3.5 minutes. Scale = 10µm. n = 3 for each genotype. (ii) Fission versus Fusion events in *foxg1a^+/+^* and *foxg1a^+/5bp^*: a fusion event can be seen as upwards inflection of the line, while fission is represented by a downwards inflection (see methods). *foxg1a^+/5bp^* embryos display increased fission events compared to *foxg1a^+/+^* siblings. Statistics were carried out as individual t-tests at each timepoint and significance can be seen as asterisks above the data point. (iii) Organelle measurements from Nellie: Organelle Area is significantly increased in *foxg1a^+/5b^* ; Organelle Tortuosity (measure of geometric complexity) shows a trend towards increase in *foxg1a^+/5bp^*; and Organelle Velocity shows no statistical difference. Statistics performed with an unpaired nested t test. (C) Bioenergetic phenotype of *foxg1a* mutants (i) Mitochondrial stress test traces for 48hpf *foxg1a^+/+^*, *foxg1a^+/5bp^* and *foxg1a^5bp/5bp^* zebrafish larvae. The x axis indicates time, and the y axis indicates oxygen consumption rate (OCR) measured in pmol/min. Control MO, n = 15 *foxg1a^+/+^*, 34 *foxg1a^+/5bp^*, and 10 *foxg1a^5bp/5bp^*. ATG4&5 MO, n = 10 *foxg1a^+/+^*, 31 *foxg1a^+/5bp^*, and 10 *foxg1a^5bp/5bp^*. Error bars indicate SDM. (ii) Basal respiration measurements for control and MO treated larvae: *foxg1a^+/5bp^* and *foxg1a^5bp /5bp^* (*p = 0.034* for *foxg1a^+/5bp^*; *p = 0.0047* for *foxg1a^5bp/5bp^*; two-way ANOVA). ATG4/5 MO injection rescues increased respiration. (iii) ATP-coupled respiration of *foxg1a^+/5bp^* and *foxg1a^5bp /5bp^* embryos injected with control or ATG4/5 MOs; *p = 0.0238* for *foxg1a^+/5bp^*; *p = 0.0004* for *foxg1a^5bp/5bp^*; two-way ANOVA). (iv) Oxidative phosphorylation (OXPHOS) of *foxg1a^+/5bp^* and *foxg1a^5bp /5bp^* embryos injected with control or ATG4/5; *p = 0.0043* for *foxg1a^+/5bp^*; *p = <0.0001* for *foxg1a^5bp/5bp^*; two-way ANOVA).

These observations led us to question whether there was a difference in mitochondrial dynamics. To answer this, we performed timelapse imaging of *foxg1a^+/+^* and *foxg1a^+/5bp^* embryos in the Tbr2;CAAX-mCherry background. Embryos were injected with mito-GFP DNA at the one cell stage to achieve mosaic mitochondrial labelling in dorsal excitatory neurons (Figure 4B). Fission versus fusion events were assessed using Nellie organelle area measurements to perform the analysis described in^23^. We found that *foxg1a^+/5bp^* had a propensity for increased fusion events (Figure 4B ii), while the *foxg1a^+/+^* siblings tended to undergo more fission events. To validate our findings from Figure 4A, we also employed Nellie to assess organelle area, tortuosity, and velocity. We again found that the *foxg1a^+/5bp^* mutant had a significant increase in organelle area (p = <0.0001, unpaired nested t-test) and a trend toward increased tortuosity (p = 0.0699, unpaired nested t-test).

Our 24hpf RNA-seq datasets (Supplementary Fig. 2) revealed that differentially expressed genes (DEGs) were enriched for mitochondrial function, including some members of the electron transport chain (ETC) e.g. *ndufa6* and *cox10*. This observation suggested a functional difference in mitochondrial output, which we then examined by performing a mitochondrial stress test on 48hpf embryos (Figure 4C). At baseline, both *foxg1a^+/5bp^* and *foxg1a^5bp/5bp^* exhibited an increase in oxygen consumption rate (OCR), compared to *foxg1a^+/+^* (Figure 4C i). Similarly, when examining ATP-coupled respiration and oxidative phosphorylation, a significant increase in OCR and ATP production rate were measured in mutants. These differences were no longer apparent in populations treated with ATG4&5 MO, demonstrating that the C-Foxg1^Mut^ directly or indirectly affects mitochondrial energy production (Figure 4C ii-v).

### C-Foxg1 interacts with ribosomal proteins and amplifies mitochondrial translation to increase neurogenesis

Given that the translation of transcripts encoded by the mitochondrial genome (mt-DNA) is the rate limiting step in the ETC, and that a proportion of Foxg1 is localised to the mitochondria, we postulated that C-Foxg1 may modulate translation within this organelle. A mass spectrometry study in mouse neuronal cells identified ribosomal proteins, including mitochondrial ribosomal proteins (Mrp), as potential Foxg1 interaction partners^24^. We selected Mrps34 (Mitochondrial ribosomal protein small subunit 34) as a candidate for further exploration as this protein is central to translation activation in the mitochondria^25^ and likely interacts with Foxg1^24^. Wholemount immunostaining of Mrps34, revealed a dramatic increase in the area and intensity of Mrps34 expression in *foxg1a^+/5bp^* and *foxg1a^5bp/5bp^* mutants at 24hpf (Figure 5A). Such differences in Mrps34 expression were not observed in the *foxg1a ^+/ ΔC-term^* and *foxg1a ^ΔC-term/ΔC-term^* mutants (Supplementary Fig. 4B iii). Complementarily, when WT *C-Foxg1-mScarlett* RNA was mosaïcly overexpressed, we observed a strong positive correlation between C-Foxg1-mScarlett and Mrps34 expression (Supplementary Fig. 4C). Together, these results demonstrate that Mrps34 protein accumulation is dependent on the C-terminal cleavage fragment. Furthermore, the colocalization of Foxg1 with Mrps34 was significantly increased in *foxg1a^+/5bp^* mutants, with this increase more apparent in the dorsal telencephalon (i.e. the location of Tbr2^+^ excitatory neurons, Figure 5A ii). Expansion microscopy of 48hpf wildtype embryos stained for Foxg1 and Mrps34 (Figure 5B) confirmed that the two peptides colocalize within the pallium.

**Figure 5:**
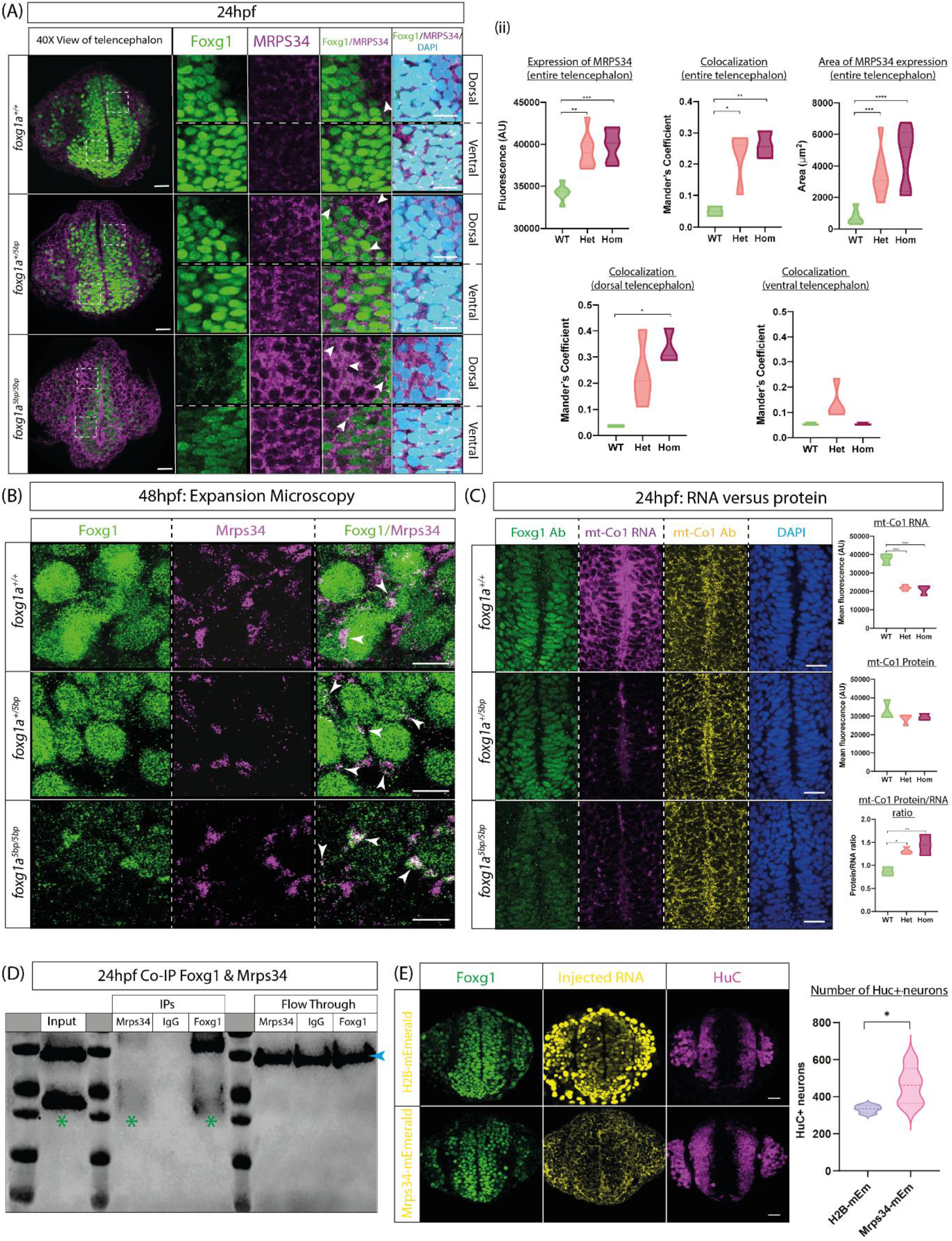
C-Foxg1 and C-Foxg1Mut interact with mitochondrial translation machinery. (A) Wholemount immunostaining of Foxg1 and Mrps34 in 24hpf zebrafish telencephalon; images show a frontal view with dorsal to the top, scale bars = 20µm (whole brain), 10µm (inset). (ii) Quantification of 24hpf Mrps34 expression and colocalization with Foxg1; *foxg1a^+/+^*(n = 3), *foxg1a^+/5bp^* (n=4) and *foxg1a^5bp /5bp^* (n=3); statistics from One-way ANOVAs. (B) Expansion microscopy of 48hpf zebrafish larvae stained for Mrps34 and Foxg1 showing colocalization of the two, scale bars = 5µm. (C) Representative images of 24hpf *foxg1a^+/+^*, *foxg1a^+/5bp^* and *foxg1a^5bp/5bp^* with mt-CoI RNA (magenta), mt-CoI protein (yellow), Foxg1 protein (green), and DAPI (blue). Plots on the right are quantification of mt-CoI RNA expression (top), protein expression (middle) and the ratio of protein/RNA (bottom). Statistics from One-way ANOVAs. (D) Co-immunoprecipitation in *foxg1a^+/+^* fish showing that the cleaved C-Foxg1 (green asterisk) interacts with mitochondrial ribosomal protein small subunit 34 (Mrps34). Gapdh (37kDa, blue arrow) shown as a loading control. Full-length Foxg1 (50kDa) is also visible in Mrps34 IP since it encompasses the C-terminal cleavage product and the IP was performed with total lysate. (E) Immunostaining of Foxg1 (green) and HuC (magenta) in 30hpf embryos injected with 100pg of either Mrps34-mEmerald or H2B-mEmerald RNA at the one cell stage. The number of HuC^+^ neurons was counted: n=4 H2B-mEmerald; n = 5 Mrps34-mEmerald, statistics from unpaired t test. Scale bar = 20µm.

To determine if mitochondrial translation is affected in *foxg1a^+/5bp^* mutants, we performed RNA fluorescent *in situ* hybridisation chain reaction (HCR) followed by immunostaining for mt-Co1 and Foxg1. Mutant telencephalons exhibited a decrease in the expression of mt-Co1 RNA, however, there was no significant difference in protein levels compared to WT (Figure 5C), indicating that translation of mt-Co1 is significantly increased in mutants. Bulk deep sequencing data from 24hpf embryos indicated that out of the 13 protein-coding genes of the mitochondrial genome, the three complex IV transcripts (mt-Co1-3) were downregulated in *foxg1a^+/5bp^* mutants (Supplementary Fig. 4D). We therefore also performed Western Blot analysis to assess expression of mt-nd1 (unchanged mRNA expression) and found an increased expression of this protein in mutant lysates (Supplementary Fig. 4D iii). To assess whether C-FOXG1 promotion of mito-translation is due to the interaction of FOXG1 with the mito-ribosome, we performed dual co-immunoprecipitation pull downs for Mrps34 and Foxg1 (Figure 5D) on protein extracts of 24hpf WT zebrafish. When probed with an antibody against Foxg1, Western Blots of the IPs showed C-Foxg1 immunoreactivity in the Mrps34 pulldown lane, demonstrating that the proteins interact in WT fish. To test whether enhanced mitochondrial translation underlies the increased excitatory neuronal population observed in *foxg1a^+/5bp^* mutants, we overexpressed Mrps34 in WT embryos and quantified neuronal numbers (Figure 5E). Overexpression of Mrps34-mEmerald, but not H2B-mEmerald RNA, resulted in an increase in the number of HuC^+^ neurons at 30hpf (p = 0.0453, unpaired t test).

### Mitochondrial function of C-FOXG1 is conserved from fish to human

Given the pathology in postmitotic neurons of zebrafish FOXG1 Syndrome models (Supplementary Fig. 4A), we performed a dual SMAD inhibition protocol on *FOXG1^256DupC+/-^* and isogenic control iPSCs (036S_CTM) to produce NPCs before terminal differentiation with DAPT treatment (notch inhibition). We initially characterised the neuronal population at day 40 *in vitro* (DIV 40) and found that *FOXG1^256DupC+/-^* samples had an increase in the number of cells present in the field of view, despite an equal number of cells being seeded for terminal differentiation (Supplementary Fig. 5B). Immunostaining for TBR1 revealed that all neurons present in the culture were excitatory, suggesting that human C-FOXG1^Mut^ also acts to increase the number of excitatory neurons. We then assessed the mitochondrial phenotype, performing immunocytochemistry for FOXG1 (C-terminal antibody) and COX-IV (Figure 6A&B). Heterozygous mutant neurons exhibited an increased mitochondrial footprint compared to isogenic controls as well as an increase in mean branch length of mitochondria (Figure 6C). When examining the subcellular distribution of FOXG1, we found a trend toward increased non-nuclear localisation in *FOXG1^256DupC+/-^* neurons, and the percentage of FOXG1 protein overlapping with the mitochondria was significantly increased. An examination of expression of MRPS34 protein revealed a significant increase in *FOXG1^256DupC+/-^* neurons compared to controls (Figure 6D). Furthermore, the percentage of non-nuclear FOXG1 protein overlapping with MRPS34 was also increased in mutant neurons (Figure 6D, iii). Our findings in human neuronal cells reflect the phenotypes identified in *foxg1a* mutant zebrafish, revealing a previously unknown mitochondrial defect in FOXG1 Syndrome and the mechanism underlying these observations.

**Figure 6:**
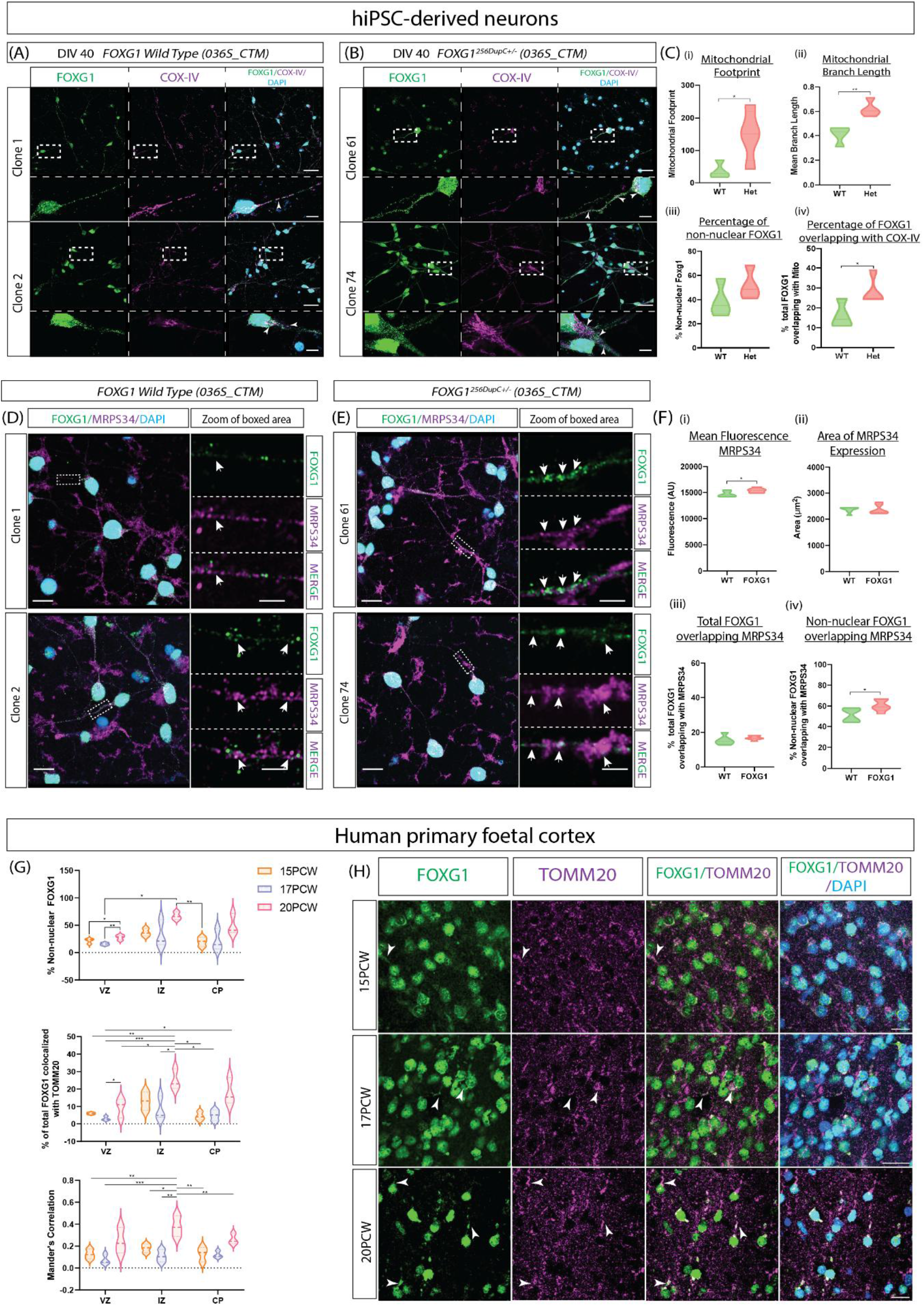
Human FOXG1 nonsense mutant iPSC-derived neuronal cells exhibit a mitochondrial phenotype. (A)&(B) Representative images from immunostaining for FOXG1 (green) and COX-IV (magenta) in DIV40 F*OXG1^256DupC+/-^* and isogenic control lines (036S_CTM). Dashed boxes indicate enlarged area in panel below. Scale bars = 20µm and 5µm (inset). n = 2 clones per genotype with 3 separate field of views for each clone. (C) Quantification of FOXG1 and COX-IV staining: all statistics from unpaired t tests. (i) Mitochondrial footprint (proxy for number of mitochondria) measured using MiNA (ii) Mitochondrial branch length (mitochondria size) (iii) Percentage of non-nuclear FOXG1 (iv) Percentage of FOXG1 overlapping with COX-IV. (D) &(E) Representative images from immunostaining for FOXG1 (green) and MRPS34 (magenta) in DIV40 isogenic control lines (D) and F*OXG1^256DupC+/-^* (E). Dashed boxes indicate enlarged area in panel to the side. Scale bars = 5µm and 2.5µm (inset). n = 2 clones per genotype with 3 separate field of views for each clone. (F) Quantification of FOXG1 and MRPS34 staining: all statistics from unpaired t tests. (i) Mean fluorescence intensity of MRPS34. (ii) Expression area of MRPS34. (iii) Percentage of total FOXG1 overlapping with MRPS34 (iv) Percentage of non-nuclear FOXG1 overlapping with MRPS34. (G) Quantification of FOXG1 colocalization with TOMM20 in primary human foetal cortex. n = 3 biological replicates for each developmental timepoint; statistics from Two-way ANOVAs. VZ; ventricular zone, IZ; intermediate zone, CP; cortical plate. (i) Percentage of non-nuclear FOXG1 (i) Percentage of FOXG1 overlapping with TOMM20. (ii) Mander’s correlation coefficient for FOXG1 and TOMM20. (H) Representative images showing FOXG1 (green) and TOMM20 (magenta) colocalization in 15, 17, and 20PCW in the human foetal IZ. Scale bars = 20µm.

To address whether the developmental dynamics of cytoplasmic and mitochondrial FOXG1 observed in zebrafish was conserved in the human telencephalon, we performed immunostaining for FOXG1 and TOMM20 in human embryonic cortical slices. The developmental stages of 15, 17, and 20 post conception weeks (PCW) were chosen due to their high rate of neurogenesis. We quantified the percentage of FOXG1 overlapping with TOMM20 and found a developmental increase, with 20PCW displaying the highest non-nuclear fraction of FOXG1 and the most colocalised to the mitochondria (Figure 6G). Further representative images showing non-nuclear FOXG1 colocalized with TOMM20 can be found in Supplementary Fig. 6C.

## Discussion

Our study reveals that Foxg1 undergoes proteolytic cleavage and localizes to mitochondria through a mechanism conserved from fish to humans, where it functions as a previously unrecognized regulator of mitochondrial translation. The cleaved C-terminal FOXG1 fragment (C-FOXG1) represents the first identified spatiotemporal regulator of mitochondrial translation in vertebrates. This newly uncovered role provides a molecular framework linking nuclear gene regulatory programs governing cell fate specification with the control of mitochondrial metabolism.

We show that severe human FOXG1 syndrome mutations (frameshift and nonsense) are not merely haplo-insufficient. Instead, patients suffer a dual molecular dysfunction: loss of full-length protein activity and gain of the cytoplasmic C-FOXG1 function. This gain arises from the use of a downstream CSC, generating a truncated protein (C-FOXG1^Mut^) which mimics the wildtype cleaved product and has a pronounced impact on the neurogenesis of excitatory pallial neurons. The availability of C-Foxg1^Mut^ from early development leads to a precocious increase in the interaction of Foxg1 with the mitochondrial translation machinery (Mrps34), resulting in a structural and functional mitochondrial phenotype. As translation is the rate limiting step in ETC activity^26^, we propose that the observed increase in the translation of mitochondrially encoded transcripts leads directly to the increase in OXPHOS, evidenced by our experiments. Repression of C-Foxg1^Mut^ expression is successful in treating excitatory cell defects, but not the loss of inhibitory cells, or the thinning of the AC. Increased Tbr2^+^ excitatory neuron numbers are therefore a consequence of C-Foxg1^Mut^ GOF, while decreased inhibitory cell numbers arise from a loss of full-length Foxg1 activity, subsequently leading to an incomplete differentiation of ventral telencephalic areas. This notion is further supported by our characterisation of the *foxg1a^+/ΔC-term^* mutant, which exhibits a decrease in both excitatory and inhibitory cell numbers at 32hpf (Supplementary Fig. 4B ii). The *foxg1a^5bp/5bp^* mutant fails to develop an AC (Supplementary Fig. 1B), with axonal bundles stalling at the lateral edge of the telencephalon. However when homozygous donor cells are transplanted into a WT background, they do indeed cross the midline, demonstrating that this is a non-cell autonomous mutant defect (Supplementary Fig. 1C) attributed to a failure of ventral specification and subsequent lack of attractive axonal guidance cues.

Genotype-phenotype correlation studies have demonstrated that early nonsense or frameshift mutations in the N-terminal of FOXG1 result in the most severe clinical outcomes, particularly with respect to seizure status^6,27^. Our findings offer a mechanistic explanation for this observation: the use of CSCs is conserved from fish to humans and overexpression of the resulting C-Foxg1^Mut^ increases excitatory neuron numbers in both species. In heterozygous mutants, this neuronal excess correlates with increased colocalization of C-Foxg1^Mut^ with mitochondria and Mrps34, most prominently in the dorsal telencephalon, and with a C-Foxg1^Mut^ dependent increase in mitochondrial size and activity. The dorsal enrichment of cytoplasmic C-Foxg1 and C-Foxg1^Mut^, combined with the selective impact of the latter on excitatory populations, suggests a dorsoventral regulation of protein localisation. This idea is supported by previous work from Hanashima et al, who showed that a DNA-binding domain-deficient form of Foxg1 was able to rescue decreased pallial cell numbers in mice lacking Foxg1^28^. Recent studies have indicated cell-type specific differences in the mitochondrial proteome, with excitatory and inhibitory cells exhibiting heterogenous expression of nuclear-encoded mitochondrial transcripts^29^. Our results expand on this concept, indicating that different cell types require a unique mitochondrial composition for their respective functions. Moreover, differentially regulated mitochondrial localisation of C-Foxg1 across the telencephalon shapes these cell type-dependent mitochondrial variations.

Along with changes in mitochondrial activity, neurogenesis is associated with dynamic alterations in mitochondria structure. Newborn neurons exhibit highly fragmented mitochondria, but more mature neurons display fused mitochondrial networks which are thought to be the structural basis underlying increased OXPHOS potential^4^. In the context of our results, the observed increase in mitochondrial size in mutant neurons suggests a more mature mitochondrial architecture compared to WT neurons (Supplementary Fig. 4A). This is also consistent with the premature differentiation phenotype that is often associated with FOXG1 syndrome and its disease models. Furthermore, antisense inhibition of C-Foxg1^Mut^ rescued the increased mitochondrial footprint, mean branch length, and functional output of the mitochondria in mutant fish, allowing us to conclude that C-Foxg1^Mut^ is responsible for these phenotypes. The proper function of the ETC is essential for efficient ATP output via OXPHOS, requiring appropriate regulation of translation within the mitochondria. Mutations in genes encoding mitochondrial translation machinery are associated with multiple neurodevelopmental phenotypes such as intellectual disability, seizures, myelination, and abnormal volume/structure of the CC ^25,30,31^. Interestingly, mitochondrial ribosomal subunits appear to be developmentally regulated with an increased expression from birth, before a peak at ∼5.5 years, which corresponds to the average age of susceptibility of neurodevelopmental disorders ^32,33^. The mechanisms that regulate mitochondrial translation remain poorly understood and to date the only mitochondria-specific translational activator identified is TACO1, which promotes ribosomal translation initiation through interaction with Mrps34^34^. C-FOXG1 is the first spatio-temporal regulator of mitochondrial translation identified, providing the means to start dissecting the mechanisms controlling mitochondrial translation at cell type resolution.

This work provides an important revision of the causes of FOXG1 Syndrome pathophysiology. Although the N-terminus of *FOXG1* is a hotspot for frameshift mutations^35^, patient variants have been reported throughout the gene and it is likely that mutations closer to the C-terminus will produce truncated proteins that lack the mitochondrial localizing domain. Such mutations are similar to our C-terminal deletion CRISPR, which exhibited a markedly different cellular E/I phenotype compared to the frameshift mutant (N-terminal lacking). Our findings also highlight a previously unrecognised therapeutic target that extends beyond FOXG1 Syndrome, as mitochondrial C-FOXG1 is likely to play a role in other disorders involving mis-regulation of FOXG1, and may serve as a useful therapeutic tool in conditions of mitochondrial hypoactivity. This work illustrates the importance of verifying the nature of early nonsense/frameshift mutations, instead of presuming these are null alleles. Nonsense mutations account for around 11% of all disease-causing mutations in humans^36^ and preliminary examination of related genes, including *MECP2* and *CDKL5*, has highlighted multiple potential CSCs downstream of reported nonsense mutations. Future work will focus of defining the roles of C-Foxg1 and C-Foxg1^Mut^ in regulating mitochondrial translation across cell types and species, and on elucidating the mechanisms underlying Foxg1 cleavage and mitochondrial import. Ultimately, as demonstrated by our phenotypic rescue experiments, such insight could lead to the development of novel antisense-based therapeutics for FOXG1 Syndrome and related human disorders.

## Acknowledgements

We would like to thank Michelle Macurak (MEH lab), who produced the (*Tg(eomesa:QF2)^c7^*^14^ *;Tg(QUAS:mApple-CAAX; he1.1:mCherry)^c636^* line; Oscar Marin for his comments and feedback on the manuscript and Graham Cocks for his production of the isogenic iPSC lines used in this study. We would also like to acknowledge Biorender.com, which was used to produce the graphic abstract for this work. This work is supported by funds from the Wellcome Trust (WT 220861/Z/20/Z to CH) and Medical Research Council (MR/N026063/1 to HB). Human Developmental Biology Initiative Wellcome (215116/Z/18/Z to KL and CH). Heart of Racing LLC Brain Health in Gen2020 to KL.

## Author Contributions

Conceptualization, HB & CH. Methodology, HB. Investigation, HB. Experimental work, HB. Mrps34-mEmerald OE experiments, KA. Human foetal tissue staining and imaging, GG. Image analysis, HB. RNA-sequencing data analysis, FH. Visualization, HB. Production of (*Tg(eomesa:QF2)^c714^ ;Tg(QUAS:mApple-CAAX; he1.1:mCherry)^c636^* line, MEH. Production of *foxg1a^+/5bp^* line, CM & OS. Mt-nd1 western blot, LOS. Production of C-terminal deletion line, HB. Writing, review and editing, HB and CH. Supervision, CH. Funding acquisition, HB, KL & CH.

## Declaration of Interests

The authors declare that they have no known competing financial interests or personal relationships that could have appeared to influence the work reported in this article.

## Materials and Methods

### Zebrafish Husbandry

All fish used in this project were cared for and raised at the Guy’s Campus Zebrafish facility,King’s College London. Embryos for experimental analysis were collected from adult zebrafish pairings and incubated at 28.5°C in fish system water containing 0.01% methylene blue; to minimise pigment formation 0.003% 1-phenyl-2-thiourea (PTU) was added to the fish water. The local Animal Care and Use Committee at King’s College London approved all fish work in this study, which was carried out under licenses issued by the Home Office and experiments were carried out in accordance with the Animals in Scientific Procedures Act of 1986. The *foxg1a* nonsense mutation used in this study was achieved by a TALEN (transcription activator-like nucleases) deletion of 5 base pairs (bp) near the start codon of the *foxg1a* coding region; this deletion results in a frameshift and the induction of an early stop codon. TALEN technology predates the recent CRISPR-Cas9 advances in genome editing and relies on the generation of a gene-specific TAL effector DNA binding domain fused to a DNA cleavage domain to mediate double stranded DNA breaks (Nemudryi et al., 2014). As the *foxg1a* homozygous mutant fish dies 8-10 days post fertilisation (dpf), adult fish stocks were maintained as heterozygous carriers. Progeny from heterozygous incrosses were genotyped by PCR amplification and subsequent sanger sequencing, genotype was determined by the deletion of 5bp.

#### Adult Zebrafish

Finclips from adult fish were digested overnight at 55°C in 50μl DNA extraction buffer (*100mM Tris-HCL pH8.5, 5mM EDTA pH 8.0, 200mM NaCl, 0.2% SDS*) supplemented with 100μg/ml Proteinase K. The following morning the Proteinase K was heat-inactivated at 85°C for 45 minutes; DNA was diluted 1:10 in molecular grade H_2_0 and 1μl was used as a template in a 25μl PCR reaction (F – GAGGAGTTGCCAGAGCAAG, R – GCCGTTTTTCTTGTCGCCTT) using DreamTaq Green PCR Mastermix (*Thermofisher*, K1081).

#### Zebrafish Larvae

Larvae up to 6dpf were genotyped by alkaline lysis DNA extraction in 45μl of solution (*25mM NaOH, 0.2mM EDTA*), heated to 95°C for one hour; 50μl of neutralisation buffer (*40mM Tris-HCL pH8.0*) was then added and 1μl was then used as DNA template in a PCR reaction.

#### Genotyping of Fixed Zebrafish Larvae

Following in situ hybridisation or wholemount immunostaining, fixed embryos were genotyped after imaging by DNA extraction from the tails of the embryo. Tails were digested overnight at 55oC in 30μl of Fixed Sample DNA Extraction Buffer (*50mM KCl, 10mM Tris-HCL pH8.3, 0.3% TWEEN-20, 0.3% NP40*) supplemented with 100μg/ml Proteinase K. The following morning the Proteinase K was heat-inactivated as before, and 2μl was used in 25μl pcr reaction. Following PCR, samples were checked on a 1% Agarose gel with ethidium bromide staining and sent for sequencing with the forward primer using Eurofins Mix2Seq Overnight Sequencing Service.

### Zebrafish Lines

The transgenic lines used in this study were obtained by crossing heterozygous *foxg1a* mutants with the established transgenics; details of each transgenic line can be found in table 1.

**Table 1:**
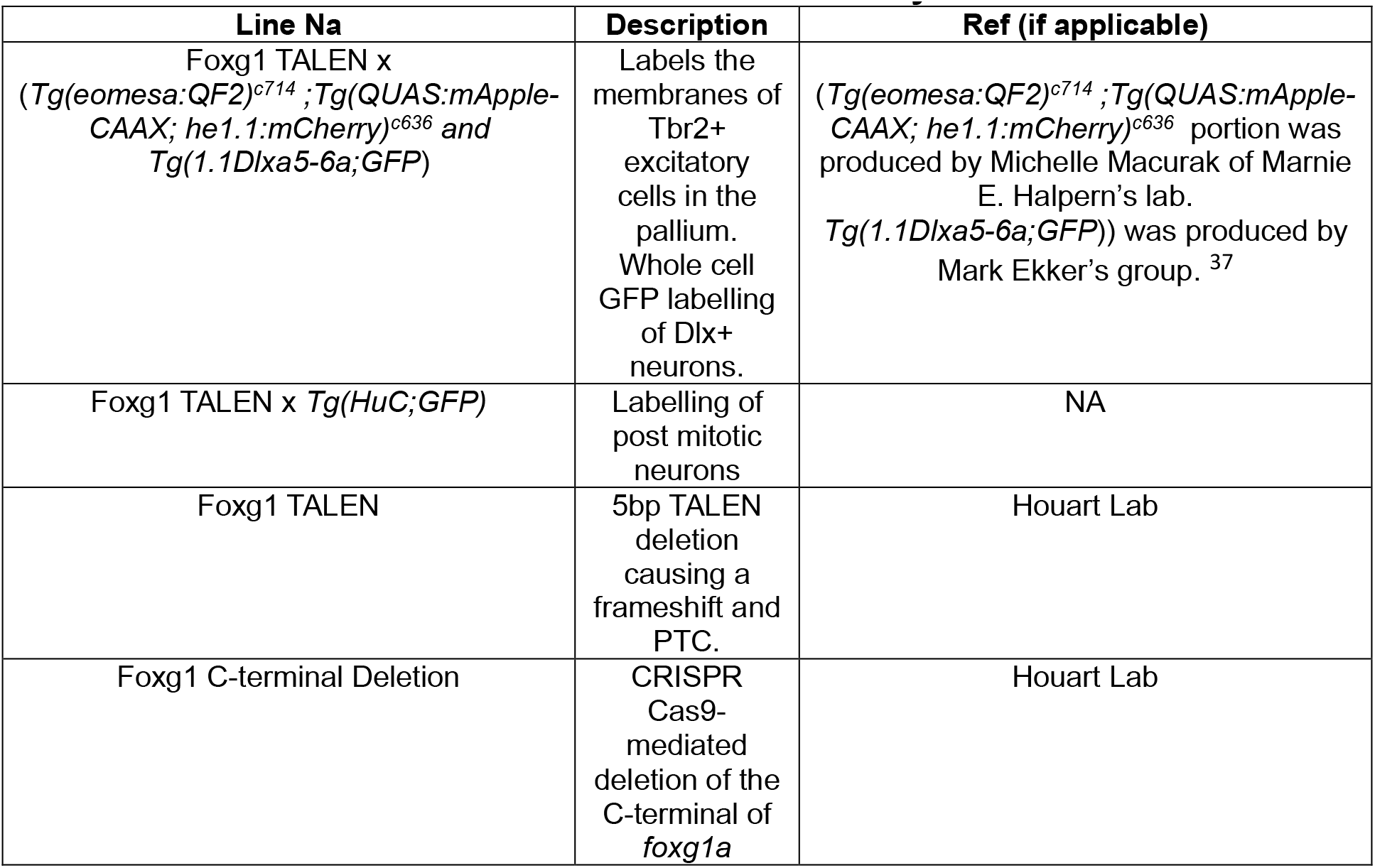
Details of zebrafish lines utilised in this study.

**Table 2:** Details of antibodies used in wholemount immunostaining of zebrafish larvae.

| Antibody | Company | Host Species | Dilution |
| --- | --- | --- | --- |
| Anti-FOXG1 (C-terminal) | Abcam, ab196868 | Rabbit | 1:500 |
| Anti-COX-IV | Abcam, ab33985 | Mouse | 1:500 |
| Anti-MRPS34 | Thermofisher, PA5-59872 | Mouse | 1:500 |
| Anti-mtCol | Abcam, ab14705 | Mouse | 1:500 |
| Anti-HuC | Abcam, ab176106 | Mouse | 1:500 |
| Anti-Rabbit Alexa Fluor 488 | Abcam, ab150077 | Goat | 1:1000 |
| Anti-mouse Alexa Fluor 568 | Abcam, ab175473 | Goat | 1:1000 |

#### Generation of C-terminal deletion CRISPR Cas9 mutant

Integrated DNA technologies (IDT) online CRSIPR sgRNA analysis was used to determine the predicted efficiency of multiple guide RNAs, and two with the best scores were ordered as crRNA (sgRNA3; *ATACACTAACTAGCATCCAC* and sgRNA4; *TCACTTACAGTCTGATCCTG*). Lyophilised crRNAs and tracrRNA were reconstituted to 200µm in IDT duplex buffer. crRNAs were annealed with tracrRNA (IDT, 1072532) by combining 1µl of each sgRNA, 1µl of tracrRNA, and 0.28µl duplex buffer in a PCR tube and heating to 95°C for 5 mins before cooling to RT. 1µl of the resultant sgRNA complexes (61µM) were combined with 1µl of Cas9 protein (IDT, 1081058) and heated at 37°C for 5 mins to produce a ribonucleoprotein complex. Embryos at the one-cell stage were then injected with 1nl of this solution diluted 1/10 in dH_2_O. Mutation frequency was checked by PCR (F; *TTTGTGGCTACCTTAGAGTCC*, R; *ATTCGGCTTGCATTGTCCGTTC*), with amplicons run on a 1% agarose gel before extracting with the Qiagen Gel Extraction Kit (Qiagen, 28704).

#### Whole Mount Immunostaining of Zebrafish Embryos

Embryos were dechorinated before fixation in 4% paraformaldehyde (PFA) – phosphate buffered saline (PBS) for 2-4 hours at room temperature (or overnight at 4°C) and washed in 0.1% Tween-PBS (PBST). Those embryos used for in situ hybridisation were transferred to 100% methanol and stored at -20°C, while those used for immunostaining were kept in PBST at 4°C until staining.

Embryos in PBST were washed twice in PBS for 10 minutes before permeabilization in 0.25% Trypsin-PBS: 24hpf, 10 min; 48hpf, 12 min; 72hpf, 16 min; 96hpf, 18 min; 120hpf 22 min. The reaction was inhibited by addition of heat inactivated goat serum (HINGS) and embryos were washed three times in PBST for 5 minutes. Blocking was carried out by incubating embryos in 10% HINGS - 0.8% Triton-PBS at room temperature for 4 hours. Primary antibodies were incubated with samples in 2% HINGS – 0.8% Triton PBS at 4°C overnight; the following day embryos were washed several times over the course of 4-6 hours before incubating with the secondary antibody at 4°C overnight. Embryos were washed in 0.8% Triton PBS for several hours before fixation in 4% PFA for 1 hour at room temperature; after washing twice in 0.8% Triton PBS embryos were transferred to 70% glycerol for long-term storage and imaging.

#### Expansion Microscopy

Immunostaining was performed as previously described before embryo heads were dissected and stored individually in PBS in PCR tubes to allow for genotyping of the tails (performed as detailed above). Following successful genotyping, dissected heads were moved to 1.5ml Eppendorf tubes containing 0.1mg/ml Acrylolyl-X SE in 0.1X PBS with one tube for multiple heads of each genotype. Expansion microscopy was then performed as previously described^38^.

#### Quantification of Phenotypic Measurements

To ensure a fair comparison, all embryos that were to be compared were imaged with the same laser power and gain using a Zeiss Axio Imager.Z2 LSM 800 Confocal.

Cell counts of Dlx^+^ inhibitory interneurons and Tbr2^+^ excitatory neurons were performed manually on confocal Z-stacks while blinded. Either the Tbr2 or Dlx channel was combined with DAPI and the counter feature on Fiji was used to mark individual cells while scrolling back and forward through the Z-stack, ensuring no cell was counted twice. Figure 1 shows an example slice with the Tbr2 channel merged with DAPI and cells counted.

Quantification of anterior commissure phenotype was also carried out manually while blinded and three measures were used to quantify the anterior commissure phenotype: thickness, width, and volume. A schematic indicating how these measurements were taken can be seen in figure 2.

To examine differences in Foxg1 protein expression in all cells of the telencephalon the “Analyse Particles” function of Fiji was used; first segmenting Foxg1 expressing cells as regions of interest (ROI) across an entire Z-stack, then measuring the individual fluorescence intensity for each ROI. When characterising cell type specific differences in Foxg1 expression mean fluorescence intensity was calculated for each individual cell type (progenitor, mid, and differentiated) by manually drawing a region of interest around each cell; the subtype was estimated based on the distance from the ventricle. A schematic showing the selection of different subtypes can be seen in figure 3; for each embryo the maximum number of cells that could fairly be discriminated was counted from two non-overlapping slices in a stack, to ensure the same cell was not included twice.

Colocalization analysis was carried out using the Fiji plugin, “Just another Colocalization Plugin” (JaCOP,), on Otsu thresholded confocal images ^39^. Alternatively, colocalization was determined by calculating the percentage of Foxg1 overlapping other stainings. Total Foxg1 area was calculated by Otsu thresholding of the raw image, maintaining the same values for each embryo and subsequent thresholding of non-nuclear Foxg1 images. The DAPI channel was used to produce a ROI and removal of this from the original Foxg1 image determined the non-nuclear Foxg1 staining. Thresholding of the COX-IV or Mrps34 channel was performed with the Otsu method to produce a second ROI and the “clear outside” function in Fiji was used to remove any Foxg1 signal outside of this ROI. Measuring this area allowed for the calculation of non-nuclear Foxg1 overlapping with COX-IV/Mrps34, while removing this ROI from the original Foxg1 image gave the percentage of total Foxg1 overlapping with these markers. The structural mitochondrial phenotype was assessed by use of the Fiji plugin “mitochondrial network analysis tool” (MiNA)^40^. For each embryo two slices from a z stack were analysed at equivalent anterior and posterior levels between samples. Mitochondrial footprint was measured across the whole telencephalon to ensure differences in cell numbers would not impact the results; for mitochondrial branch length manual ROIs were selected based on the anatomical position.

#### Quantification of neuron versus progenitor numbers

Antibody staining for Foxg1 was performed on mutant and WT fish in a HuC;GFP background line at 3dpf and 10dpf. Quantification was performed on an anterior and posterior slice from a Z-stack spanning the entire telencephalon, as defined by the extent of Foxg1 staining. Otsu thresholding of the HuC signal was used to generate a mask that allowed for the differentiation of Foxg1+ neurons (inside HuC mask) and progenitors (outside HuC mask). Manual cell counts were performed and the average number of progenitors and neurons (across anterior and posterior slices) for each embryo were plotted. Fiji’s analyse particles feature was used to measure the mean fluorescence of Foxg1 in individual cells, with the mean value for each embryo plotted.

### RNA-sequencing

Single embryos from a *foxg1a^+/-^* in-cross were dechorinated at the appropriate timepoint (8-somite stage or 24 hours). Embryos were homogenised in buffer RLT from the RNeasy Micro Kit (Qiagen*, 74004*) using a plastic pestle, and RNA extraction was performed as per the manufacturer’s instructions, with the inclusion of the optional on-column DNase digestion. The concentration of RNA was determined using the Quibit^TM^ RNA High Sensitivity Assay (Thermofisher*, Q32852*) and the quality determined by sending an aliquot to the Bioanalyzer facility at King’s Genomic Centre, Waterloo Campus. RNA samples with an RNA integrity number (RIN) of less than 8 were discarded and those with the highest RNA concentrations were sent to the genomics facility at the Huntsman Cancer Institute High-Throughput Genomics Facility, University of Utah, USA. Library preparation was carried out using the Illumina TruSeq Total RNA Library Prep Ribo-Zero Gold kit which included depletion of ribosomal RNA prior to cDNA synthesis. Libraries were then sequenced by application to a Illumina NovaSeq flow cell on a NovaSeq 6000 instrument; each sample was read with 2x 50bp paired-end sequencing, with an average of 30 million reads per sample.

All bioinformatics was carried out by Fursham Hamid, a Postdoctoral researcher in the lab at the time of these experiments. In brief, resulting FASTQ files from the RNA-seq experiment were subjected to initial quality control steps before aligning to the Genome Reference Consortium Zebrafish build 10 genome, (GRCz10), with 70-80% of reads aligning to the reference genome. Differential gene expression analysis was performed using Whippet and an excel file containing gene names, log fold changes, and corrected significance values for each genotype were supplied to me along with BAM files. I was then able to characterise changes in gene expression using log fold changes and manual inspection of sequencing reads in IGV to determine which genes may be promising candidates for further investigation. RNA-sequencing data is available online at ArrayExpress under the accession number E-MTAB-13886.

### Whole Mount In Situ Hybridisation

A whole embryo cDNA library was produced for use as a template for in situ probe PCR by extraction of RNA from 24 hpf embryos using the Qiagen RNeasy Mini Kit (*Qiagen,* 74104), with inclusion of the optional on-column DNase digest. RNA was reverse transcribed using the First Strand cDNA Synthesis kit (*Thermo Scientific*, K1612), starting with 1μg of RNA and using Oligo(dt) primers. The concentration of resulting cDNA was measured using a nanodrop and diluted to 50ng/μl for use as a template in the production of gene-specific anti-sense RNA probes. Gene-specific mRNA primers were designed using ensemble sequences, with the reverse primer modified to contain a T3 RNA polymerase promoter sequence that would allow the transcription of an anti-sense RNA probe (primers can be seen in table x). PCR reactions for amplification of the gene-specific cDNA templates comprised 50ng of total cDNA, a final concentration of 0.5μM of each of the forward and reverse primers, and 12.5μl Dream Taq Green PCR Master Mix 2X (*Thermo Scientific*, K1081) in a total reaction volume of 25μl; cycling conditions can be seen in table 3.

**Table 3:**
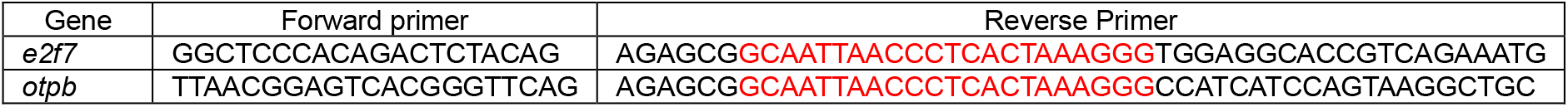
Gene-specific primers for production of *in situ* hybridisation probes. Red indicates the T3 RNA polymerase site for *in vitro* transcription.

Following amplification, gene-specific PCR products were purified with the MinElute PCR Purification kit (Qiagen, 28004) and 500ng was subsequently used in an in vitro transcription reaction. In a total volume of 20μl, 2μl of digoxgenin-labelled (DIG) NTPs (*Roche,* 11277073910), 2μl 10X transcription buffer (*Roche*), 1μl Pyrophopatase (Thermo Scientific, EF0221), 1μl RNase Inhibitor (*Roche,* 10877200), and 1.5μl of T3 RNA polymerase (*Roch,* 11031163001) were added to PCR templates and incubated at 37oC for 1 hour, 1.5μl of DNase was then added and reaction incubated for a further 15 minutes.

DIG labelled RNA probes were then purified using Mini Quick Spin Columns (*Roche,* 11814427001) before addition of 1 volume of saline-sodium citrate (SCC) and 2 volumes of formamide to the total RNA.

Embryos collected as previously described were rehydrated from 100% methanol to PBST gradually, at room temperature, before permeabilising for 10 – 30 minutes with Proteinase K in PBST (1:1000 dilution). After washing twice in PBST, embryos were fixed in 4% PFA for 20 minutes at room temperature and washed again in PBST. 100μl Hybridisation mix (*10% TWEEN, 50% formamide, 5X SCC, 0.5mg/ml Torula mRNA, 0.05mg/ml heparin*) heated to 65°C was added to the embryos, shaking at room temperature until they settled; embryos were moved to 65°C and Hybridisation mix replaced before incubating for 3-6 hours. Anti-sense RNA probes were diluted accordingly in pre-warmed Hybridisation mix, applied to the embryos and hybridised overnight at 65°C.

Embryos were washed in 65°C Hyb mix for 30mins, in 2X SSE (*3M Sodium chloride, 0.3M sodium citrate, pH 7*) 0.1% CHAPS multiple times over 1.5 hours and then several times in 0.2X SSE 0.1% CHAPS over 1.5 hours before washing in 0.1% TWEEN Maleic acid buffer (MABT) for 1 hour. Blocking was performed in MAB/blocking buffer for 2 hours before adding MAB/Anti-DIG (1:2000) AP Fab fragments (*Roche*, 11093274910) and incubating overnight at 4°C. Embryos were washed over the course of 6 hours in MABT followed by multiple washes in TBST, leaving overnight at 4°C.

Embryos were washed twice in NTMT (*0.1M NaCl, 0.1M Tris-HCl, 0.05M MgCl, 0.1% TWEEN*) pH9 for 5 minutes before incubation in 3.375μl NBT (*Roche,* 11383213001) and 3.5μl BCIP (*Roche,* 1183221001) per 1.5ml NTMT. Staining was monitored and colouration ended when reaching the desired intensity before washing several times in PBST. Following a final fixation step in 4% PFA for 1 hour at RT embryos were washed in PBST and stored in 70% glycerol until imaging, using a Nikon eclipse E800 microscope.

### Wholemount RNA Hybridisation Chain Reaction (HCR)

HCR was performed as described in (35), followed by immunostaining as described previously in this manuscript, omitting permeabilization with trypsin and using PBS-0.1% TWEEN. Quantification of mRNA versus protein expression was performed using Fiji’s analyze particles function on Otsu-thresholded images and the mean fluorescence and area of staining was calculated.

### Protein Extraction from Zebrafish Embryos

Protein for Western Blot experiments was extracted from 24hpf zebrafish embryos obtained from an in cross of *foxg1a^+/-^, Dlx4-6;GFP* adults. Use of the Dlx transgenic background enabled genotyping of embryos by eye under epifluorescence microscopy: *foxg1a^5bp/5bp^* fish exhibit an obvious lack of Dlx+ interneurons in the telencephalon and *foxg1a^+/5bp^* have a significant decrease that is visible to the trained eye. Previous validation of this genotyping method gave a 95% success rate for validating heterozygous mutants, while homozygous mutants were always correctly identified.

Yolks were removed from dechorinated embryos by washing them twice in Ringer’s Solution (*116mM NaCl, 2.9mM KCl, 1.8mM CaCl_2_, 5mM HEPES pH 7.2*), before adding Ringer’s solution complete with Protease Inhibitor Cocktail (*Roche, 11836170001*) and tirating with a pulled glass pipette. Next, embryos were centrifuged for 3 minutes at 1000g and the supernatant removed, fresh Ringer’s solution (without Protease Inhibitor Cocktail) was added to the samples and the centrifugation carried out again; this wash step was repeated three times. NP40 buffer, complete with Protease Inhibitor Cocktail (*Roche,* 11836170001), was added to the samples and incubated on ice for 10 minutes before homogenising with a sterile syringe. Samples were then centrifuged at 4°C at maximum speed for minutes to remove the insoluble cellular material, with the supernatant formed after this step being transferred to a new Eppendorf.

Protein concentration was quantified using the Pierce^TM^ BCA Protein Assay (*Thermofisher,* 23225), as per the manufacturer’s instructions. Protein samples were then either directly used in SDS page or stored at -80°C until needed.

### Immunoprecipitation (IP)

IP was performed with the Pierce^TM^ Crosslink IP (*Thermofisher, 26147*) kit using total protein lysate extracted at 24hpf as previously described. Antibodies were bound to agarose resin as detailed in the user manual by incubating with rotation at 4°C overnight prior to crosslinking with DSS (included in kit) for 45mins at RT. Pre-clearing of total lysate was performed as indicated using 1mg of total lysate for each IP (*Mrps34; PA5-59872, IgG; 02-6102, and Foxg1; ab196868*). Captured peptides were eluted with low pH buffer before denaturation at 95°C for 10mins.

### Zebrafish Western Blot

Protein lysate was diluted in NuPAGE^TM^ LDS Sample Buffer (*Thermofisher,* NP0007) and heated to 70°C for 10 minutes. 15μg of total protein lysate was run per well of a NuPAGE^TM^ 10% Bis-Tris Mini Protein Gel (*Thermofisher,* NW00105BOX), alongside a Dual Precision Protein Standard (*BioRad,* 1610374), for 1 hour at 100V. Protein was then transferred to a membrane using the iBlot2 machine, with iBlot2 PVDF Regular Stacks (*Thermofisher,* 1B24001), and template programme P0 (20V, 1 minute; 23V 4 minutes; 25V 2 minutes). The blot membrane was then blocked in 5% Milk – TBST for 45 minutes at room temperature. After blocking, primary antibodies were added in 2% Milk – TBST overnight at 4°C while shaking (*Rabbit anti-C-terminal FOXG1* ab196868*, 1:500; Rabbit anti-N-terminal FOXG1* PAB30085*, 1:500; Rabbit anti-GAPDH, 1:500, Rabbit anti-mt-nd1 ab181848, 1:500*). The following day the blot membrane was washed 3 times in TBST, before adding the secondary antibody, goat anti-rabbit IgG HRP (*Abcam,* ab6721) in 2% Milk-TBST and incubating at room temperature for 1 hour. The membrane was then washed three times with TBST before incubation with Pierce^TM^ ECL Plus Western Blotting Substrate (*Thermofisher,* 32132), as per the manufacturer’s instructions. Blot membranes were exposed using LICOR Odyssey XF imaging system.

Quantification of the relative expression of different C-terminal Foxg1 fragments was carried out in Fiji using the “analyse gels” feature. Each biological replicate consists of pooled embryos from a single clutch, with different replicates collected at different times. These were used to calculate relative expression differences normalising to the housekeeping gene GAPDH as outlined in ^42^

### Timelapse imaging of mitochondrial dynamics

Embryos from a *foxg1^+/5bp^* in cross in the Tbr2:mCherry background were injected with mito-GFP DNA at the one cell stage to achieve mosaic mitochondrial labelling in dorsal excitatory neurons. At 28hpf embryos were sorted based on the presence of Tbr2:mCherry and mito-GFP signal in the same cells, with homozygous mutants discounted. Larvae were anaesthetised and mounted in a frontal view in 0.8% low melting point agarose in fish water. Once the agarose set the petri dish was flooded with fish water without methylene blue and regions of interest imaged using airyscan confocal microscopy with a 20X water dipping objective and 5X zoom. Regions of interest were selected based on the presence of Tbr2+ neurons (mCherry) and the GFP-channel alone was used to acquire z stacks every 2.8 seconds. Analysis of fission/fusion events were performed using Nellie’s automatic organelle area measurement to measure the total organelle area at each timepoint^43^. To assess dynamic changes in organelle area we took the area at timepoint 0 and divided it by itself to obtain a starting value of 1, for each subsequent timepoint we divided the area by the previous timepoint area (e.g. t2-t1; t3-t2 etc) to determine fission/fusion events, with an increased value relative to the previous indicating more fusion events.

### Cloning and Constructs

All restriction enzyme digests were carried out using the NEB cloner online protocol database (https://nebcloner.neb.com/#!/). Following digestion all products were PCR purified with Qiagen’s MinElute PCR purification kit (28004) and DNA concentrations measured using a nanodrop instrument. Ligations were performed using the Instant Sticky-end Ligase Master Mix (*NEB*, M0370S), as per the manufacturer’s instructions. Molar ratio of ligation mixes ranged from 1:3 – 1:7.

Transformation of cloned vectors was achieved by incubating 2μl of ligation mix, or 10ng of vector, with NEB 5-alpha Competent E.Coli (C2987H) on ice for 30 minutes before heat shock at 42°C for 45 seconds and returning to ice for 2 minutes. Outgrowth was allowed in SOC Medium while shaking at 37°C for 45 minutes before plating on an appropriate antibiotic selective agar plate and incubating at 37°C overnight.

Single colonies were selected, grown overnight in antibiotic selective agar broth, and plasmid purified using Qiagen’s MiniPrep Spin Kit (*Qiagen*, 27104). DNA concentration was measured using a nanodrop instrument and vectors sent for sequencing to confirm correct cloning.

### QUAS: Syp-GFP

For telencephalic-specific labelling of excitatory synapses, a 5’ UAS: Synaptophysin-GFP vector (gift from Meyer lab) was modified to include a QUAS binding sequence instead of the classic Gal4;UAS system. This allowed for microinjection of the QUAS: Syp-GFP construct into our Foxg1 TALEN x Tg(tbr2:QF2)c714;Tg(QUAS:mApple-caax:HE:mCherry) zebrafish line, resulting in the labelling of only excitatory synapses. The UAS sequence from the original vector was excised by PciI (*NEB*, R0655S) and HindIII (*NEB*, R0104S) restriction enzyme double digest. Primers were designed against the QUAS sequence in the transgene of Tg(tbr2:QF2)c714;Tg(QUAS:mApple-caax:HE:mCherry), including a PciI restriction digest site in the forward primer (5’-TTGCTCACATGTCAACTTTGTATAGAAAAGTTGG-3’) and a HindIII site in the reverse (5’-AGCAAAAAGCTTCTTTTTTGTACAAACTTGACGC-3’). Genomic DNA from was extracted as previously described and a 230bp product was amplified using the DreamTaq Green PCR Mastermix (*Thermofisher*, K1081). The PCR product was then PCR purified with Qiagen’s MinElute PCR purification kit (28004) and digested with PciI and HindIII. The cut vector and the digested PCR product were then re-purified, ligated and transformed as described above.

Following conformation of cloning success by Sanger sequencing, 60pg of QUAS:Syp-GFP was injected into Tg(tbr2:QF2)c714;Tg(QUAS:mApple-caax:HE:mCherry) in the *foxg1a* background. Embryos were then fixed and imaged as previously described and the Fiji plugin, “SynQuant” was used to characterise synaptic phenotypes. The SynQuant plugin allowed for automated detection of synaptic puncta based on QUAS:Syp-GFP images; an example of puncta detection can be seen in figure 4^44^.

### Morpholino Design Against Cryptic ATGs

The size of the truncated protein present in the *foxg1a* homozygous and heterozygous mutants was determined using Western Blot; the size in bp was then determined using a kDa to bp calculator and the *foxg1a* gene was inspected for potential cryptic start sites. Two ATGs were identified that could produce a protein of ∼35kDa (figure 5), since they were in direct sequence with each other one MO was designed to target them both. From this point on these ATGs are denoted ATG_4 & ATG_5; this nomenclature was determined by counting the in-frame ATGs along the *foxg1a* gene. MOs were designed by Gene Tools Inc using submitted sequences with the target ATG highlighted: MO sequence against ATG_4 & ATG_5: GGCCATCATAATCAAAGCGTTGTAG. The MO was diluted to 0.6mM and 1nl injected into 1-cell stage embryos, before either extracting protein for Western Blot or fixation for antibody staining. MO acts to inhibit translation from ATG start sites and does not degrade the target mRNA.

### Mitochondrial Stress Test of 48hpf Zebrafish Larvae

Embryos from a *foxg1a^+/5bp^* incross were injected with 1nl of 0.6mM of either a control MO or the MO against ATG4&5. Embryos were dechorinated at 24hpf and maintained at 28°C in pH 7.4 E3 media until they reached 48hpf. The Seahorse flux cartridge (*103792-100, Agilent)* was rehydrated with 200μl of sterile water per well overnight at 37°C and the Seahorse XF machine was calibrated to room temperature. Drug stocks were prepared in pH7.4 E3 media such that the final concentration upon injection of 20μl per well was as follows: Oligomycin; 25μm (*495455-10MG, Sigma*), FCCP; 8μm (*C2920-10MG, Sigma*), Antimycin A; 1.5μm (*A8674-50MG, Sigma*), and Rotenone; 1.5μm (*R8875-10MG, Sigma*). Following overnight rehydration, the sterile water was removed from the Seahorse cartridge and replaced with 200μl per well of calibrant solution pre-warmed to 37°C (*100840-000, Agilent*). For each well 20μl of prepared drug solution was loaded into the corresponding injection port (Oligomycin; A, FCCP; B, Antimycin A and Rotenone; C). The sensor cartridge was then loaded into the machine and calibration commenced. Embryos for the Mitochondrial Stress Test were moved to a new dish containing pH7.4 E3 media supplemented with MS222 and a P1000 pipette set to 180μl was used to transfer individual larvae to each well of the test cartridge, leaving the four corner wells devoid of fish to allow for background readings. After calibration was completed the first cartridge was removed and replaced with the cartridge containing anesthetised larvae before beginning the Mitochondrial Stress Test Cycling, as detailed in table 4. Following completion of the procedure, individual larvae were genotyped as previously described and data was analysed using the online Wave software from Agilent.

**Table 4:** Cycling conditions for Seahorse Mitochondrial Stress Test.

|  | mix | wait | measure | cycles |
| --- | --- | --- | --- | --- |
| Baseline | 2 | 1 | 2 | 12 |
| Oligomycin | 2 | 1 | 2 | 20 |
| FCCP | 2 | 1 | 2 | 8 |
| Rotenone/AA | 2 | 1 | 2 | 12 |

### Cell Culture

#### iPSC Culture

Isogenic iPSC lines were produced by Graham Cocks of the King’s College London Genome editing core facility and obtained as liquid nitrogen frozen stocks. Alternatively, FOXG1 Syndrome patient iPSCs containing a *FOXG1 256DupC^+/-^* mutation were obtained from Angus Clarke at Cardiff University as frozen stocks. Cell lines were rapidly thawed and resuspended in StemFlex^TM^ media (*Thermofisher, A3349401*), supplemented with 10μm ROCK inhibitor. Cells were plated on Nunc^TM^ treated 6-well cell culture plates (*Thermofisher, 140685*) previously coated in Geltrex^TM^ (*Thermofisher, A1413301*), as per the manufacturer’s instructions. After 24 hours media was replaced with StemFlex^TM^ media without ROCK inhibitor, following which, media was replaced every 2 days. iPSCs were cultured until 70-80% confluency before passaging at a ratio of 1:6: cells were washed with room temperature (RT) Hanks Balanced Salt Solution (HBSS; *Thermofisher; 14170088*) and dissociated by adding 1ml Versene (*Thermofisher, 12569069*) incubated for 3-5mins at 37°C. Versene was aspirated from the culture vessel and StemFlex^TM^ media was used to collect the cells gently with a cell scraper. Frozen cell stocks were prepared by dissociating cells as described and resuspending iPSCs from a single well of a 6-well plate in 1ml of StemFlex^TM^ 10% DMSO (*Sigma, D2650*). Cells were cooled to -80°C at a rate of -1°C per min using a Mr Frosty container (*Thermofisher, 5100-0001*), after 24 hours cells were transferred on dry ice to liquid nitrogen for long term storage.

#### Neuralisation

iPSCs for neural induction were cultured on Geltrex coated plates until 100% confluent to ensure generation of CNS-specific neuronal precursor cells (NPCs). When confluency was reached StemFlex^TM^ media was aspirated, and Neuralisation media (components in table 5) was added to the cells - the addition of neuralisation media was denoted as day 1 in the neuralisation timeline. Addition of Dorsomorphin and SB43 to culture media allowed for neural induction via dual SMAD inhibition: SB43 acting on activin/nodal/TGF-β and Dorsomorphin inhibiting both activin/nodal/TGF-β and BMP pathways ^45^.

**Table 5:** Components of neuralisation media for dual SMAD inhibition protocol.

| Media | Component |
| --- | --- |
| N2 | 49ml DMEM (*Sigma*, D6421), 0.5ml 100x N2 (*ThermoFisher*, 1750248), 0.5ml 100X Glutamax (*ThermoFisher*, 35050-061) |
| B27 | 48.5ml Neurobasal (*ThermoFisher*, 21103049), 1ml 50X B27 (*ThermoFisher*, 17504044), 0.5ml 100X Glutamax (*ThermoFisher*, 35050-061) |
| Neuralisation Media (N2B27) | 50% N2, 50% B27, 1μM dorsomorphin (*Sigma*, P5449-5MG), 10μM SB43 (*Sigma*, S4317-5MG) |

Neuralisation media was replaced every 24 hours until day 8, when NPCs were passaged at a ratio of 1:1. After aspiration of media, cells were washed in RT HBSS before the addition of 1ml Accutase (*Sigma, A6964-500ML*) for each well of a 6-well plate. NPCs were incubated at 37°C for 3-5mins until able to be lifted from the vessel with gentle pipetting, cells were transferred to a falcon tube containing RT DMEM and centrifuged for 2 mins at 900RPM. The supernatant was aspirated, cells resuspended in RT DMEM and centrifuged for a further 2 mins to ensure all Accutase had been eliminated. NPCs were then resuspended in Neuralisation media without the addition of Dorsomorphin and SB43, supplemented with 10μm ROCK inhibitor to promote cell survival and plated on Geltrex coated plates. Media was replaced every day until day 13, when cells were once again passaged at a ratio of 1:1; on day 16 and 23 cells were passaged as described, but at a ratio of 1:2.

For terminal differentiation of neurons, coverslips were coated with Poly-D-Lysine (*A3890401, Thermofisher*) diluted to 50μg/ml in Borate buffer (*28341, Thermofisher*) and incubated at 37°C for at least 4 hours. Coverslips were then washed three times with PBS before replacing with laminin (*Merck, L2020-1MG*) diluted to 9.6μg/ml in cold DMEM and incubated overnight at 37°C. NPCs for terminal differentiation were passaged with Accutase, as previously described, and plated on laminin coated coverslips at a density of 200,000 cells/cm^2^ in B27 media supplemented with 10µM DAPT (*Stem Cell Technologies, 72082*), 200µM AA2P (*Merck, A4403*), and 10µM ROCKi. Following 24 hours of incubation the total volume of media was removed and replaced with B27 supplemented with DAPT and AA2P; media was replaced every day for 7 days following terminal plating. Following this, a part media change was performed every 7 days omitting DAPT from the media. Coverslips for immunocytochemistry were fixed in 4% PFA for 10 mins at RT before permeabilization in 2% HINGS 0.4% TRITON-PBS for 30 mins. Primary antibodies added in 2% HINGS 0.4% TRITON-PBS and incubated overnight at 4oC before washing 3X in 0.4% TRITON-PBS, secondary antibodies were incubated for 1 hour at RT in 2% HINGS 0.4% TRITON-PBS and samples washed 3X before mounting. Details of antibodies for ICC can be found in table 6.

**Table 6:** Antibodies used in human ICC.

| Antibody | Company | Host Species | Dilution |
| --- | --- | --- | --- |
| Anti-FOXG1 (C-terminal) | Abcam, ab196868 | Rabbit | 1:100 |
| Anti-COX-IV | Sigma, T6793 | Mouse | 1:100 |
| Anti-MRPS34 | Thermofisher, PA5-59872 | Mouse | 1:100 |
| Anti-NeuN | Abcam, ab104224 | Mouse | 1:100 |
| Anti-Tbr1 | Abcam, ab183032 | Rabbit | 1:100 |
| Anti-Rabbit Alexa Fluor 488 | Abcam, ab150077 | Goat | 1:1000 |
| Anti-mouse Alexa Fluor 568 | Abcam, ab175473 | Goat | 1:1000 |

### Human tissue

Human foetal neocortical material was provided by the Joint MRC/Wellcome Trust (grant# MR/X008304/1 and 226202/Z/22/Z) Human Developmental Biology Resource (HDBR; http://hdbr.org). Three biological replicates from post-conception weeks (pcw) 15, 17, and 20 were analyzed. Upon arrival, tissues were fixed in 4% (wt/vol) paraformaldehyde (PFA) at 4 °C for at least 24h, incubated in 15% sucrose for 24h, and subsequently stored in 30% sucrose until processing.

### Tissue processing and imaging

Foetal neocortical tissue was immersed and incubated in 7.5% porcine skin gelatine (Sigma-Aldrich, G1890) in 15% sucrose for 4 h at 37°C in a water bath and kept overnight at 4 °C. The next day samples were snap-frozen in 2-methylbutane (Sigma-Aldrich, 277258) at −60 °C and stored at −80 °C until sectioning. Cryosections were obtained at 20 µm and 150 µm thickness. Sections of 150 µm were used for expansion microscopy, performed as previously described, while 20 µm sections were used for immunofluorescence.

For immunofluorescence, sections were stained with primary antibodies against FOXG1 (1:100, Abcam ab196868), TOMM20 (1:100, Santa Cruz sc-17764), and MAP2 (1:100, Antibodies A85363). Secondary antibodies were applied at 1:1000 dilution: donkey anti-rabbit Alexa Fluor 488 (Abcam ab150073), donkey anti-mouse Alexa Fluor 568 (Thermo Fisher A10037), and donkey anti-chicken Alexa Fluor 647 (Invitrogen A78952). DAPI (Sigma D9542) was used for nuclear counterstaining. All antibodies were diluted in 10% donkey serum containing 0.3% Triton X-100. Sections were mounted in Mowiol (Merck Biosciences).

Images were acquired using a Nikon Ti2 AX inverted confocal microscope. Sections were imaged with a 20×/0.8 NA air objective and 60×/1.42 NA oil immersion objective. A 1.5× intermediate magnification was applied in all acquisitions.

### Analysis of FOXG1 and TOMM20 colocalization

Fiji was used as previously described to quantify the percentage of FOXG1 overlapping with TOMM20, while the plugin JaCoP was used to calculate Mander’s correlation coefficients.

## Abbreviations

Corpus Callosum; CC, Autism Spectrum Disorder; ASD, Interneurons; IN, Excitatory/Inhibitory; E/I, Premature Termination Codon; PTC, Cryptic Start Codon; CSC, Anterior Commissure; AC, Neuronal Progenitor Cells; NPCs, Morpholino; MO, Electron Transport Chain; ETC, VZ; Ventricular zone; SVZ; subventricular zone, IZ; intermediate zone, SP; subplate, CP; Cortical plate.

---

**Figure 1:**
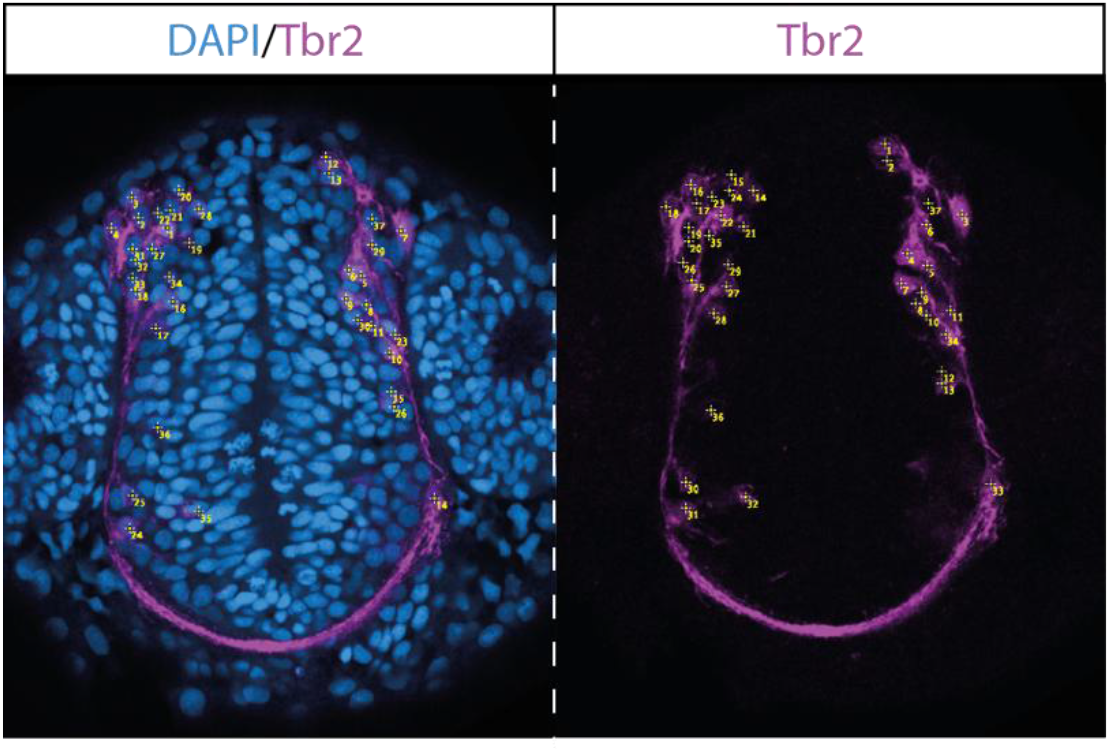
Example of Tbr2 cell counting in a 28hpf embryo. Either Tbr2 (shown) or Dlx channel was combined with DAPI and cells counted with the counting feature in Fiji by scrolling back and forward through a Z-stack.

**Figure 2:**
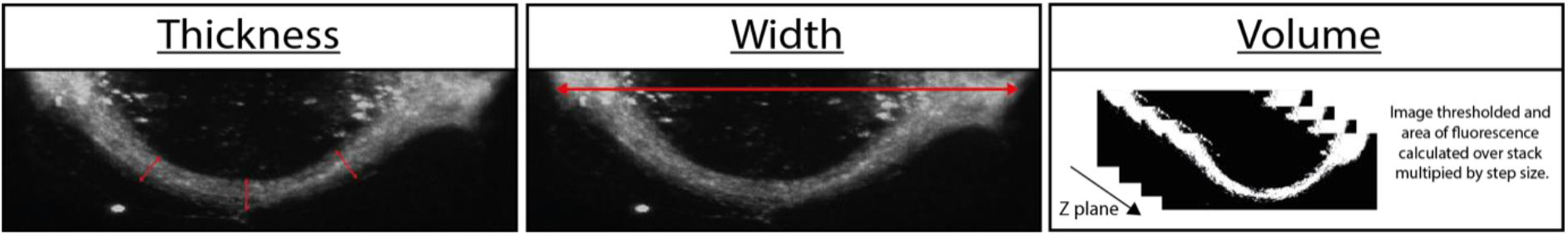
Phenotype measures used in the quantification of anterior commissure (AC) phenotypes. The thickness of the AC was measured in three locations (indicated in figure) following which an average value was calculated for each sample. Width was defined as the length from each lateral edge of the AC. Volume was measured by calculating the area of the AC across the entire 3D-projected image and multiplying by the step size.

**Figure 3:**
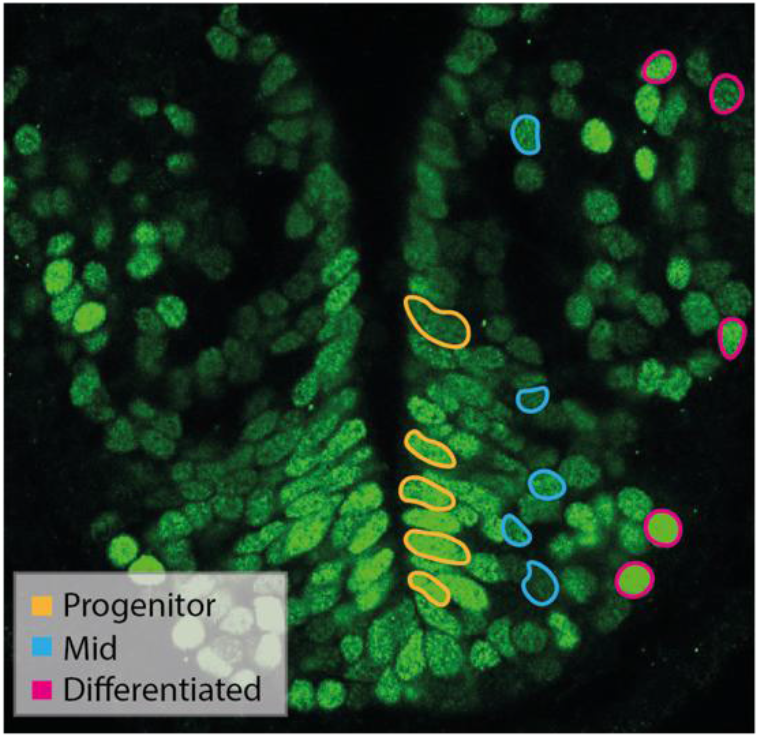
Selection of cells for cell type specific measurements of Foxg1 expression at 2dpf. Cell identity was determined based on location relative to the ventricular wall. Progenitors were situated directly adjacent to the wall (yellow); migrating/differentiating cells (referred to as “Mid”) were assigned to those cells between the progenitor population and the cells at the most lateral edges of the telencephalon; postmitotic neurons (“Differentiated”) were assigned to those at the lateral-most edges of the telencephalon.

**Figure 4:**
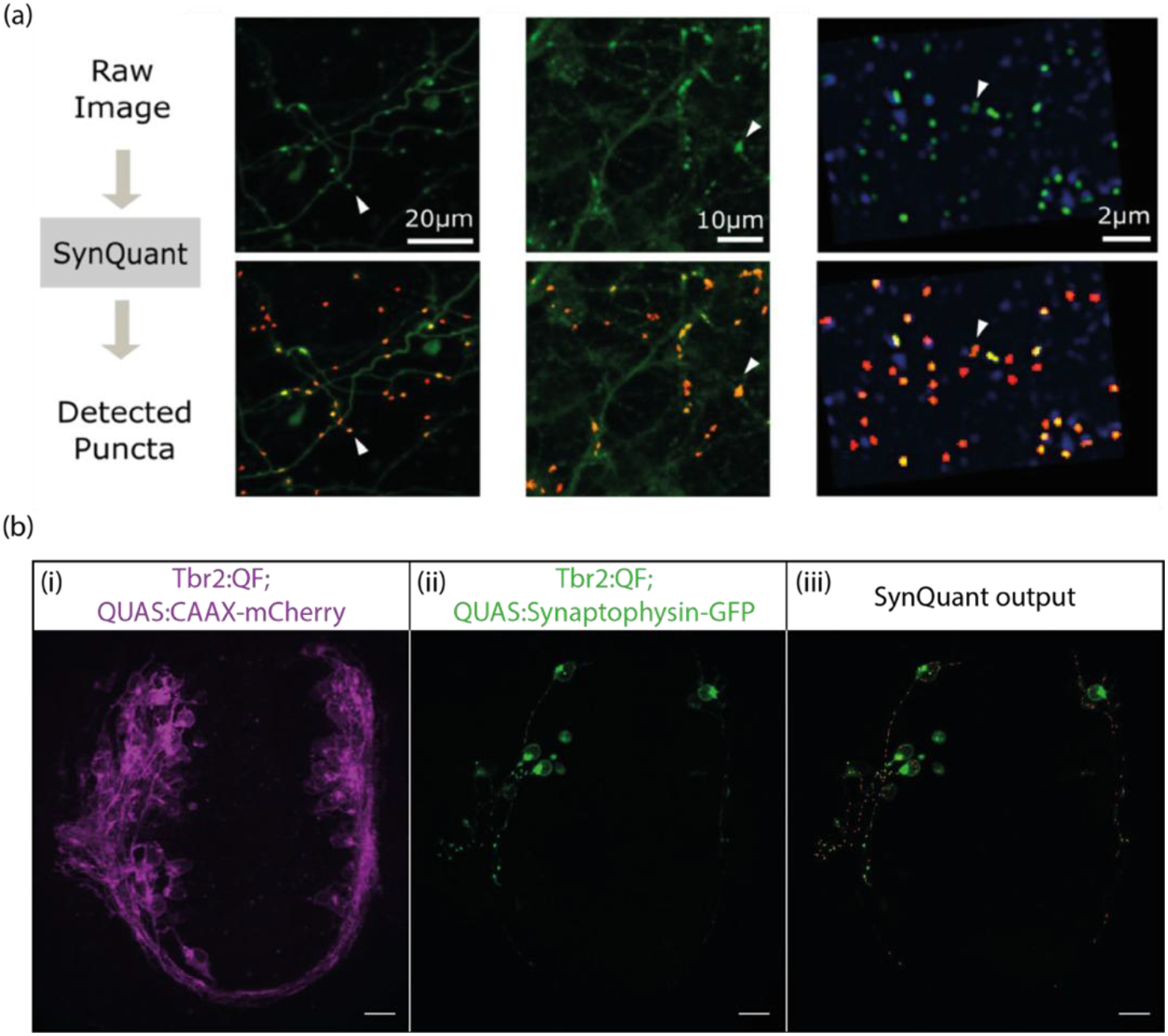
Automatic detection of synaptic puncta using SynQuant. **(a)** Examples of input images and the resultant detected synaptic puncta from Wang et al, 2020. **(b)** Representative images from labelling of excitatory synapses in *foxg1a* mutants using SynQuant. **(i)** Confocal projection of Tbr2+ cell membrane labelling in our transgenic line. **(ii)** Example of labelling achieved upon injection of QUAS:Synaptophysin-GFP construct, showing that labelling colocalises only with Tbr2+ cells. **(iii)** Representative SynQuant output for the automatic detection of synaptic puncta on QUAS:Synaptophysin-GFP images.

**Figure 5:**
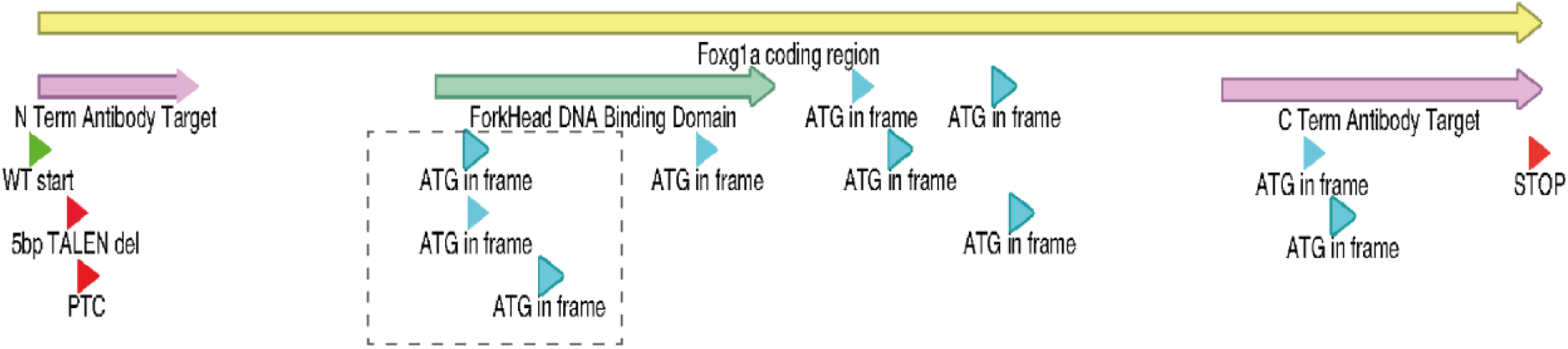
Schematic of the *foxg1a* coding region. The position of the 5bp TALEN deletion and subsequent premature termination codon (PTC) are indicated in red. Downstream in-frame ATGs are indicated in blue, with the three potential cryptic start codons (CSCs) in use highlighted by a dashed box.

## Notes

https://www.ebi.ac.uk/biostudies/arrayexpress/studies/E-MTAB-13886?query=E-MTAB-13886

### Competing Interest Statement

The authors have declared no competing interest.

### Summary of Updates

We have added characterisation of a C-terminal deletion CRISPR mutant to confirm the selectivity of the mitochondrial phenotype to the PTC model. Further characterisation of mitochondrial phenotypes has been assessed with timelaspe imaging in zebrafish, while we also show FOXG1 colocalises with mitochondria in primary human fetal brain samples.

