## Supplementary Figures for "A cleaved cytosolic FOXG1 promotes excitatory neurogenesis by enhancing mitochondrial translation"

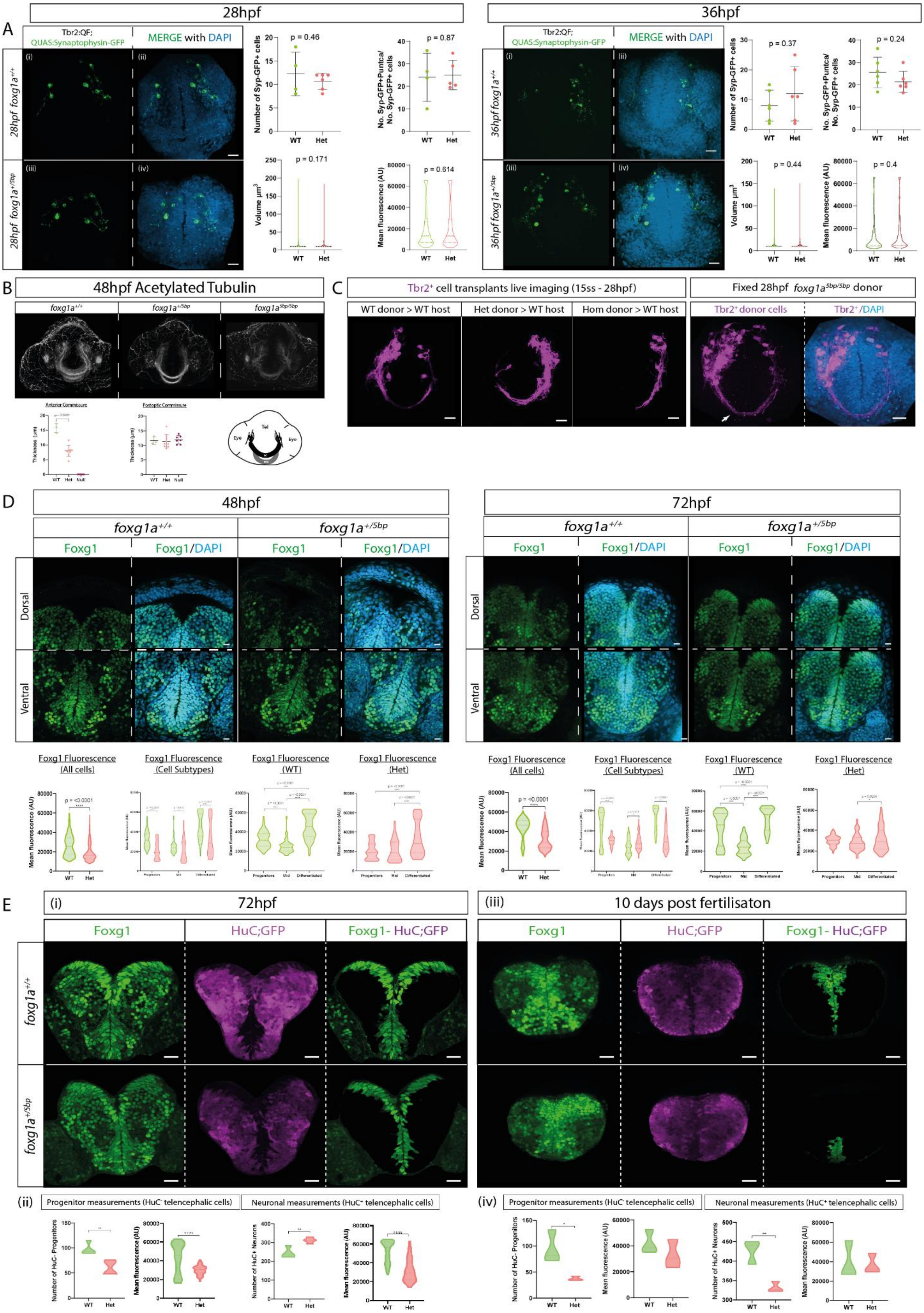

### Supplementary Figure 1: extended phenotypic characterisation of *foxg1a*$^{+/5bp}$ mutant.

A. Synaptophysin synapse labelling in Tbr2+ postmitotic neurons: QUAS:Synaptophysin-GFP DNA injection into Tbr2:QF embryos at one cell stage for mosaic excitatory synapse labelling. At 24hpf and 36hpf there was no significant difference between *foxg1a*$^{+/+}$ and *foxg1a*$^{+/5bp}$ fish in any of the measurements performed (number of synaptophysin puncta normalized to number of positive cells, volume of puncta, and mean fluorescence of puncta, which were assessed by unpaired t tests). 24hpf: n = 4 *foxg1a*$^{+/+}$, 6 *foxg1a*$^{+/5bp}$. 36hpf: n = 4 *foxg1a*$^{+/+}$, 6 *foxg1a*$^{+/5bp}$. Scale bars = 20μm; error bars indicate SEM.

B. Antibody staining for acetylated tubulin, with the *foxg1a*$^{+/5bp}$ fish exhibiting a significant decrease in the thickness off this commissure at 48hpf. The homozygous mutant did not have an anterior commissure in any of the individuals examined. Measurement of the postoptic commissure thickness (present outside the limits of Foxg1 expression), which was not significantly altered in either the heterozygous or homozygous mutant. n = 3 *foxg1a*$^{+/+}$, 9 *foxg1a*$^{+/5bp}$, 7 *foxg1a*$^{5bp/5bp}$; Scale bars = 20μm; error bars indicate SEM.

C. Cell transplantation experiments: Tbr2$^{+}$ cells were transplanted from either *foxg1a*$^{+/+}$, *foxg1a*$^{+/5bp}$, or *foxg1a*$^{5bp/5bp}$ donors at ~50% epiboly to WT hosts and principal neuron development imaged overnight from the 15ss. *foxg1a*$^{5bp/5bp}$ donor cells were able to extend axonal processes and cross the midline.

D. Analysis of Foxg1 protein expression at 48 and 72hpf. The mean fluorescence of Foxg1 staining was significantly decreased in the heterozygous fish at 48hpf (p = <0.0001, unpaired t test). The expression of Foxg1 was significantly decreased in all three cell subtypes in *foxg1a*$^{+/5bp}$ fish at 48hpf (p = <0.0001 for progenitors, p = 0.013 for Mid cells, p = <0.0001 for neurons; unpaired t tests). In WT fish the expression of Foxg1 significantly decreased between progenitor and Mid cells (p = <0.0001; one way ANOVA), while neurons had the highest expression of Foxg1. The decreased expression of Foxg1 in Mid cells was not observed in the heterozygous mutant, but like the WT, neurons had a significantly increased expression of Foxg1 (p=<0.0001; One way ANOVA). At 72hpf Foxg1 expression was again significantly decreased in all cells in the heterozygous telencephalon (p = <0.0001; unpaired t test). Comparison of cell subtype expression between WT and heterozygous mutants at 72hpf indicated that Foxg1 was lower in expression in progenitors (p = <0.0001; unpaired t test), higher in expression in Mid cells (p = 0.0071; unpaired t test), and lower in expression in neurons (p = <0.0001; unpaired t test) in the *foxg1a*$^{+/5bp}$ mutant. The expression pattern of Foxg1 between cell subtypes followed the same trend at 72hpf, with *foxg1a*$^{+/5bp}$ fish failing to show a decreased expression in Mid cells while the difference in neuronal Foxg1 expression was less pronounced. 48hpf: n = 6 *foxg1a*$^{+/+}$, 7 *foxg1a*$^{+/5bp}$; 72hpf: n = 3 *foxg1a*$^{+/+}$, 4 *foxg1a*$^{+/5bp}$. Scale bars = 20μm; error bars indicate SEM.

E. Representative images are single slices of a confocal stack taken in a frontal view; dorsal to the top a. At 3 days post fertilization (dpf) the heterozygous mutant exhibited a significant decrease in the number of HuC$^{-}$ progenitors (p = 0.0085, unpaired t test) and the expression of Foxg1 (mean fluorescence) was significantly decreased in progenitors (p = <0.0001). The number of HuC$^{+}$ neurons was significantly increased at 3dpf (p = 0.0053, unpaired t test) and the expression of Foxg1 was significantly decreased in this population of cells (p = <0.0001, unpaired t test) n = 4 *foxg1a*$^{+/+}$, 4 *foxg1a*$^{+/5bp}$. By 10dpf the number of HuC$^{-}$ progenitors significantly decreased in *foxg1a*$^{+/5bp}$ fish (p = 0.036, unpaired t test), with no difference in the expression of Foxg1. The number of HuC$^{+}$ neurons was significantly decreased in *foxg1a*$^{+/-}$ (p = 0.0083, unpaired t test), however, the expression of Foxg1 was no longer increased. n = 3 *foxg1a*$^{+/+}$, 3 *foxg1a*$^{+/5bp}$. Scale bars = 20μm; error bars indicate SDM.

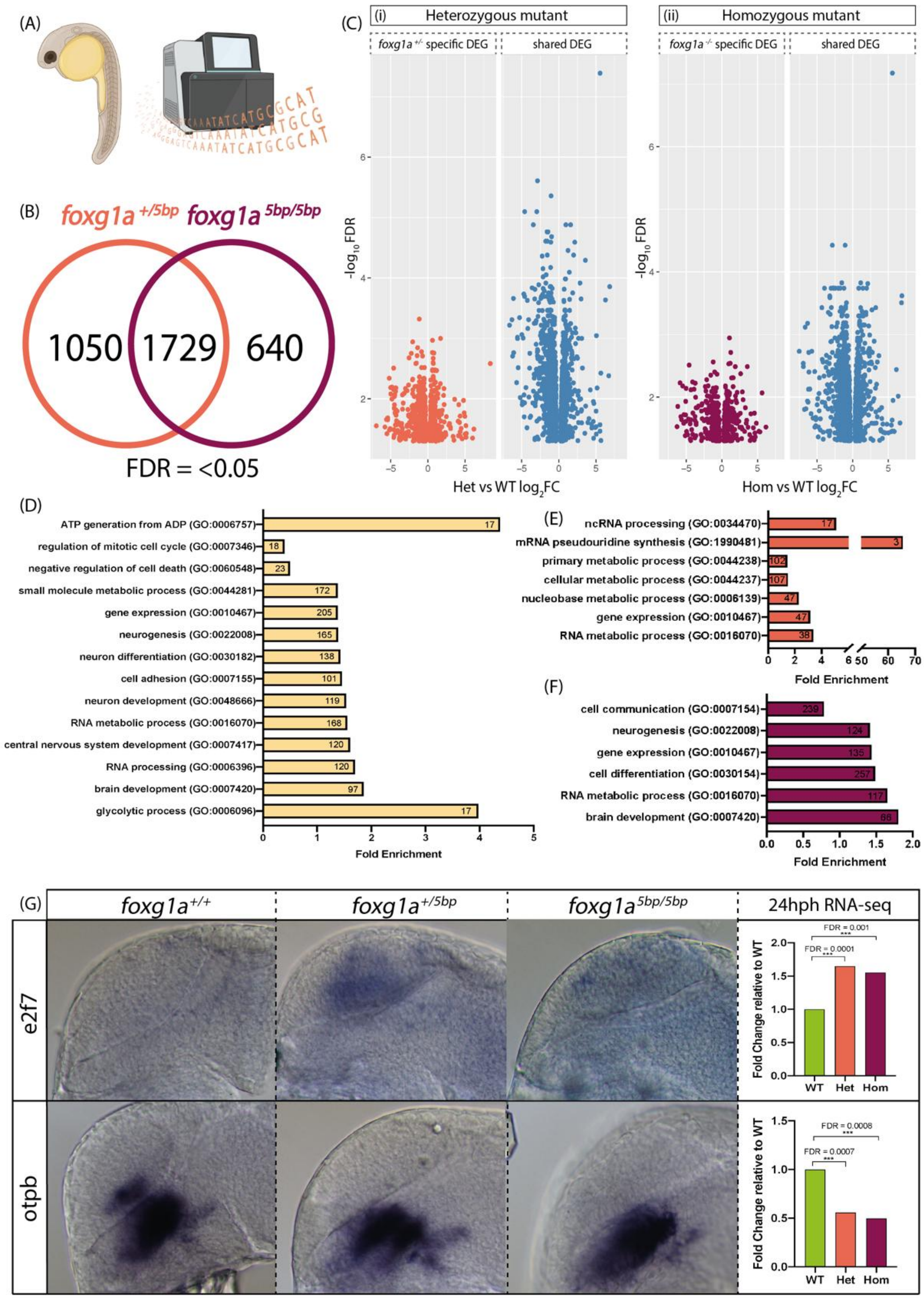


### Supplementary Figure 2: Molecular phenotype of the heterozygous *foxg1a* nonsense mutant at 24hpf.

A. 24hpf RNA-sequencing: n = 5 *foxg1a*$^{+/+}$, 7 *foxg1a*$^{+/5bp}$, 4 *foxg1a*$^{5bp/5bp}$.

B. Differential gene expression analysis revealed a total of 3419 differentially expressed genes (DEGs) compared to *foxg1a*$^{+/+}$; 1729 of these were shared between *foxg1a*$^{+/5bp}$ and *foxg1a*$^{5bp/5bp}$ embryos; 1050 were specifically changed in *foxg1a*$^{+/5bp}$ and 640 were specific to *foxg1a*$^{-/-}$ (false discovery rate (FDR) = <0.05).

C. Volcano plots showing the significant DEGs in the heterozygous and homozygous mutant: those genes specifically changed in each mutant are shown on the left in coral (*foxg1a*$^{+/5bp}$) or purple (*foxg1*$^{5bp/5bp}$), and genes which are also differentially expressed in both mutants are shown on the right in blue.

D. Selected significant Panther Gene Ontology (GO) terms that were enriched in all DEG: x axis denotes fold enrichment within the gene set and the numbers on bars indicate the number of genes in each GO classification.

E. Selected significant Panther GO terms that were specifically enriched in *foxg1a*$^{+/5bp}$ mutants.

F. Selected significant Panther GO terms that were specifically enriched in *foxg1a*$^{5bp/5bp}$ mutants.

G. In situ hybridization as validation of 24hpf RNA-seq: *e2f7* shown as an example of an upregulated gene and *otpb* as an example of a downregulated gene. Graphs to the right indicate the fold change values from the RNA-seq.

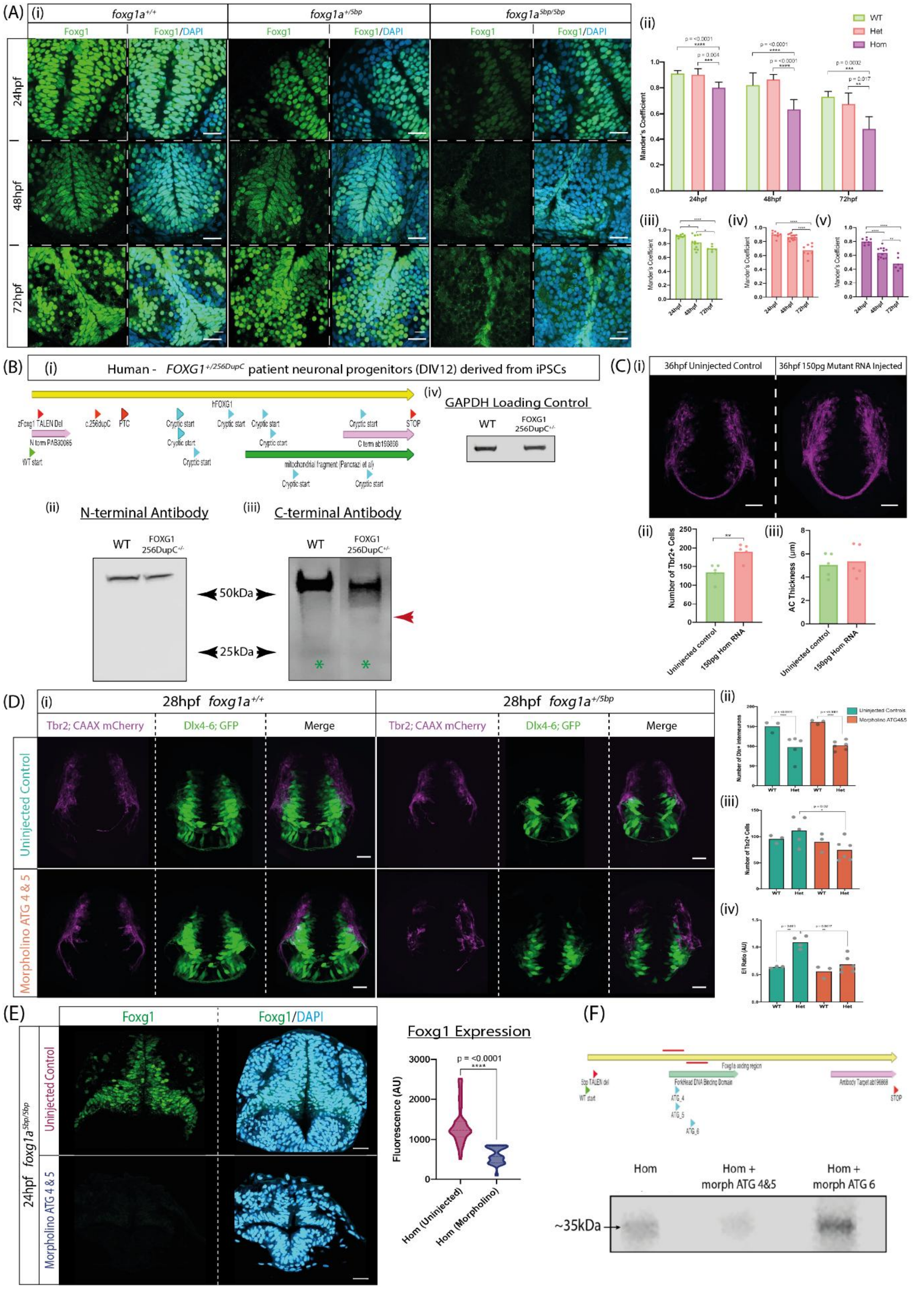

**Supplementary Figure 3: induction of a premature stop codon does not prevent translation.**

(A) Foxg1 staining with antibody raised against the C-terminus of the protein.

(i) Representative images of 24hpf, 48hpf, and 72hpf embryos of each genotype stained for Foxg1 and DAPI in a frontal view, scale bars = 20μm.
(ii) Quantification of Foxg1 and DAPI colocalisation at 24hpf, 48hpf, and 72hpf; Mander's coefficient of 1 equates to perfect colocalisation; One-way ANOVA.
(iii) Quantification of Foxg1 and DAPI colocalisation for *foxg1a*$^{+/+}$; One-way ANOVA.
(iv) Quantification of Foxg1 and DAPI colocalisation for *foxg1a*$^{+/5bp}$; One-way ANOVA.
(v) Quantification of Foxg1 and DAPI colocalisation for *foxg1a*$^{5bp/5bp}$; One-way ANOVA.

(B) Western Blot experiments repeated on DIV 12 neuronal progenitors derived from FOXG1 Syndrome patient iPSCs. The patient carries the FOXG1256DupC$^{+/-}$ mutation present in our isogenic iPSC lines. FOXG1 C-terminal antibody blot also detects a smaller peptide of ~35kDa, indicating that the cryptic start codon (CSC) is used in patient cells to produce C-FOXG1$^{Mut}$ (indicated by red arrow).

(C) Overexpression of mutant mRNA transcript at a later timepoint of 36hpf.
(i) Representative images showing frontal view of uninjected control and 150pg mutant RNA injected WT fish.
(ii) Quantification of number of Tbr2$^{+}$ cells; injected embryos exhibit a significant increase (p = 0.0048, unpaired t test).
(iii) Quantification of anterior commissure thickness; no significant difference (unpaired t test).

(D) Morpholino treatment of *foxg1a*$^{+/+}$ and *foxg1a*$^{+/5bp}$ at an early timepoint of 28hpf.
(i) Representative images of 28hpf embryos in a frontal view, scale bars = 20μm.
(ii) Number of Dlx$^{+}$ interneurons; Two-way ANOVA.
(iii) Number of Tbr2+ excitatory cells; Two-way ANOVA.
(iv) E/I cellular ratio; Two-way ANOVA.

(E) Validation of morpholino against cryptic start codons ATG4&5 (CSCs). Representative images of homozygous nonsense mutants stained for Foxg1 following injection of morpholino against CSCs. Quantification of Foxg1 expression shows a significant decreased upon treatment with morpholino (p = <0.0001; unpaired t test).
(F) Western blot for Foxg1 (C-terminal Ab) on *foxg1a*$^{5bp/5bp}$ mutants injected with either: control morpholino (MO; first lane), MO against ATG4&5 (suspected CSC), MO against ATG6.

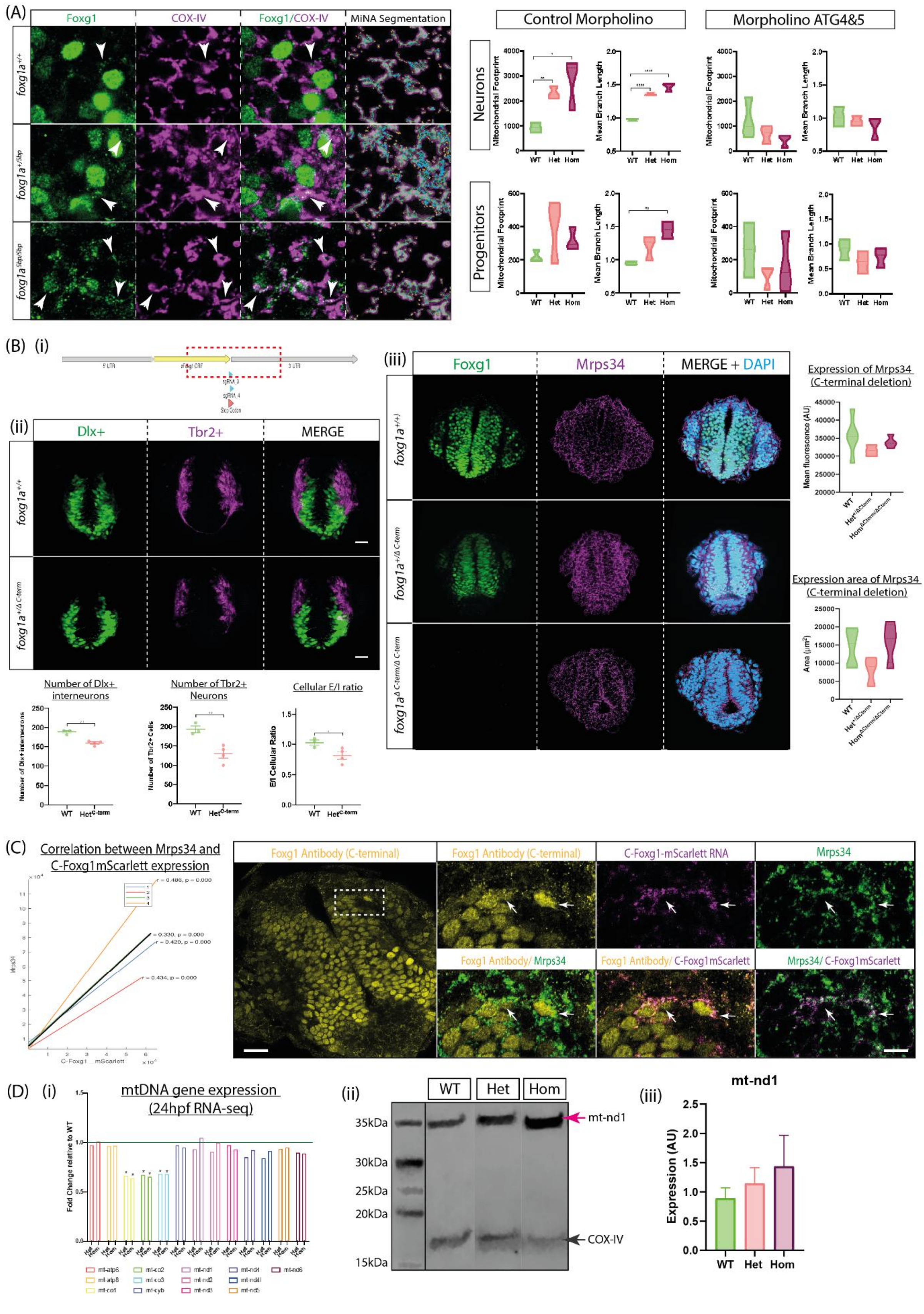


**Supplementary Figure 4: Role of WT C-Foxg1 and further characterisation of the mitochondrial phenotype.**

(A) Panel showing MiNA segmentation of mitochondria for quantification in Figure 3A. Quantification of mitochondrial phenotype in neurons and mitochondria following MO treatment. *foxg1a* mutants exhibit an increased mitochondrial footprint and mean branch length in neuronal cells, while the progenitor phenotype is less pronounced. Treatment with MO ATG4&5 rescues neuronal and progenitor mitochondrial phenotype. Control MO n = 3 *foxg1a*$^{+/+}$, 4 *foxg1a*$^{+/5bp}$, 3 *foxg1a*$^{5bp/5bp}$; ATG4&5 MO n = 3 *foxg1a*$^{+/+}$, 3 *foxg1a*$^{+/5bp}$, 3 *foxg1a*$^{5bp/5bp}$.

(B) C-terminal *foxg1a* gene deletion CRISPR.

i. Schematic of *foxg1a* gene showing positions of sgRNAs used in production of a C-terminal deletion mutant. The deletion encompasses the entire C-terminal and 3' UTR.

ii. Characterisation of E/I phenotype at 32hpf in C-terminal *foxg1a* gene deletion CRISPR. *foxg1a*$^{+/\Delta C\text{-}term}$ mutants exhibit decreased Dlx+ interneuron numbers, decreased Tbr2+ numbers, and a decrease in cellular E/I ratio. (n = 4 *foxg1a*$^{+/+}$, 4 *foxg1a*$^{+/\Delta C\text{-}term}$; statistics from unpaired t test).

iii. Antibody staining for Foxg1 (green) and Mrps34 (magenta) at 24hpf. Note lack of signal in Foxg1 antibody staining for *foxg1a*$^{\Delta C\text{-}term/\Delta C\text{-}term}$ mutants as the antibody targets the C-terminus. Scale bar = 20µm. Quantification of mean fluorescence of Mrps34 expression showing no difference between genotypes One way ANOVA (n = 5 *foxg1a*$^{+/+}$, 4 *foxg1a*$^{+/\Delta C\text{-}term}$, & 3 *foxg1a*$^{\Delta C\text{-}term/\Delta C\text{-}term}$). Quantification of area of Mrps34 fluorescence showing no difference between genotypes One way ANOVA (n = 5 *foxg1a*$^{+/+}$, 4 *foxg1a*$^{+/\Delta C\text{-}term}$, & 3 *foxg1a*$^{\Delta C\text{-}term/\Delta C\text{-}term}$).

(C) Mosaic overexpression of *C-Foxg1-mScarlett* RNA (WT cleavage fragment; magenta) followed by antibody staining for Foxg1 (yellow) & Mrps34 (green). Scale bar = 20µm (overview); Scale bar = 10µm (inset). (ii) Correlation of C-Foxg1-mScarlett fluorescence intensity with Mrps34 fluorescence intensity shows a highly significant positive correlation. Coloured lines on chart indicate single embryos; black line is line of best fit.

(D) Expression of mt-DNA genes.

i. RNA expression of the 13 protein coding genes of the mitochondria genome from bulk RNA-sequencing at 24hpf.

ii. Western blot on protein lysates from *foxg1a*$^{+/+}$, *foxg1a*$^{+/5bp}$, & *foxg1a*$^{5bp/5bp}$ probed for mt-nd1 (36kDa) and COX-IV (16kDa).

iii. Quantification of mt-nd1 expression, normalized to COX-IV expression; n= 3 per genotype.

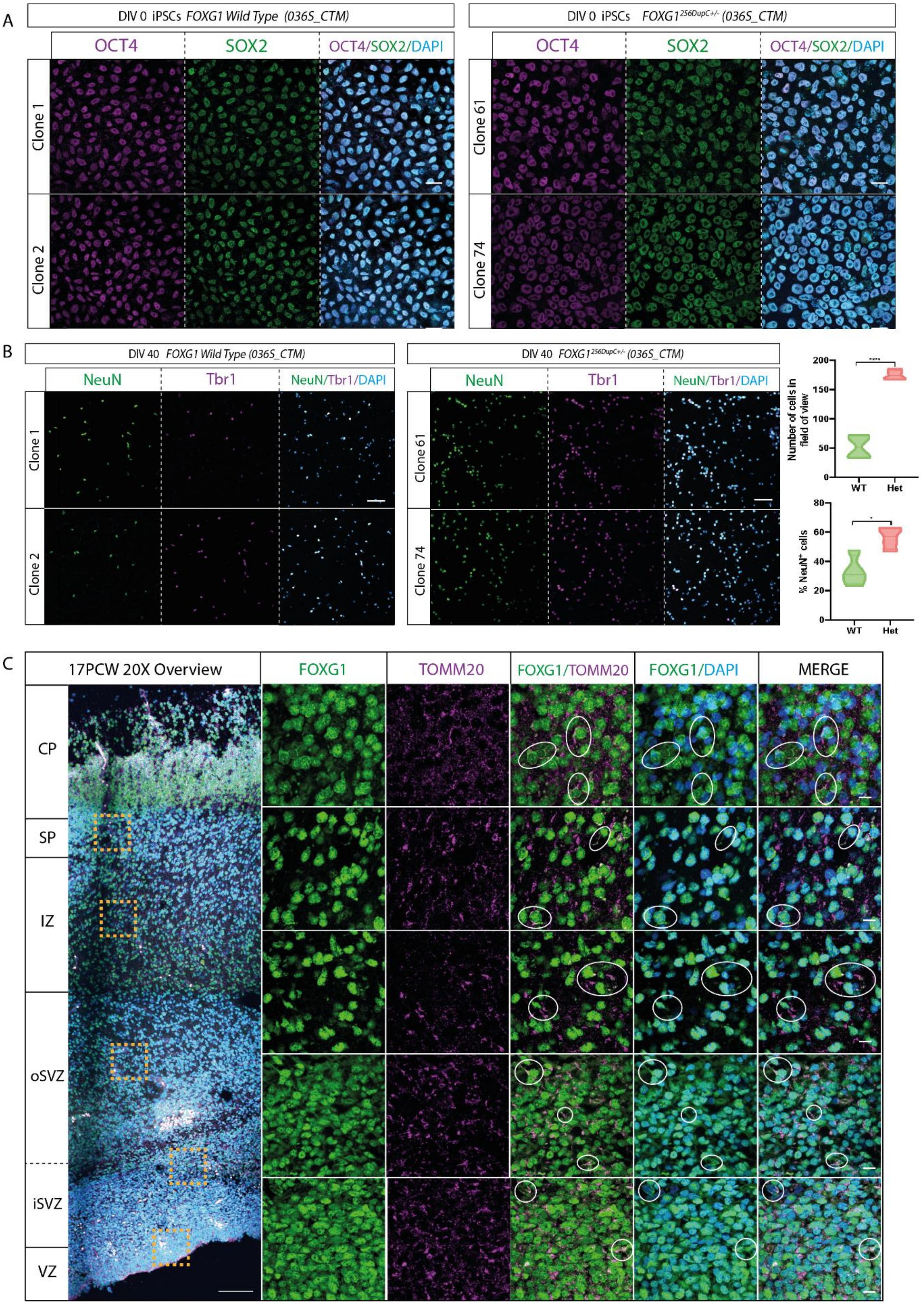


**Supplementary Figure 5: Human iPSC/Neuronal characterisation.**

(A) Immunostaining of iPSCs for OCT4 (Magenta) and SOX2 (Green), confirming pluripotency of 036S_CTM clones and *FOXG1*$^{256DupC+/-}$ isogenic clones. Scale bars = 20µm.

(B) Immunostaining of *DIV40* neuronal cells for NeuN (Green) and Tbr1 (Magenta), Scale bars = 50µm.

Quantification of number of cells in the field of view, showing *FOXG1*$^{256DupC+/-}$ samples have significantly more cells (unpaired t test, p = <0.0001). n = 2 clones per genotype, with two fields of view per clone.

Quantification of the percentage of NeuN$^{+}$ cells (post mitotic neuronal marker). *FOXG1*$^{256DupC+/-}$ samples have a significant increase in the percentage of NeuN+ cells (unpaired t test, p = 0.012). n = 2 clones per genotype, with two fields of view per clone.

(C) Representative immunostaining of 17PCW primary human foetal telencephalon: FOXG1 (green), TOMM20 (magenta), and DAPI (blue). CP; cortical plate, SP; subplate, IZ; Intermediate zone, oSVZ; outer subventricular zone, iSVZ; inner SVZ, VZ; ventricular zone. Scale bars = 100µm (overview), 10µm (insets). Circled ROIs indicate examples of FOXG1 and TOMM20 colocalization.
